# Post-fertilization transcription initiation in an ancestral LTR retrotransposon drives lineage-specific genomic imprinting of *ZDBF2*

**DOI:** 10.1101/2023.10.30.564869

**Authors:** Hisato Kobayashi, Tatsushi Igaki, Soichiro Kumamoto, Keisuke Tanaka, Tomoya Takashima, So I. Nagaoka, Shunsuke Suzuki, Masaaki Hayashi, Marilyn B. Renfree, Manabu Kawahara, Shun Saito, Toshihiro Kobayashi, Hiroshi Nagashima, Hitomi Matsunari, Kazuaki Nakano, Ayuko Uchikura, Hiroshi Kiyonari, Mari Kaneko, Hiroo Imai, Kazuhiko Nakabayashi, Matthew C. Lorincz, Kazuki Kurimoto

## Abstract

The imprinted *ZDBF2* gene is controlled by oocyte-derived DNA methylation, but its epigenetic regulation is quite different from that of other canonically imprinted genes that are dependent on DNA methylation deposited in the gametes. At the *ZDBF2* locus, maternal DNA methylation in the imprinted differentially methylated region (DMR) does not persist after implantation. Instead, a transient transcript expressed in the early embryo exclusively from the unmethylated paternal allele of the DMR, known as *GPR1-AS* in humans and *Liz* in mice, contributes to establishing secondary DMRs that maintain paternal expression of *ZDBF2* in the somatic lineage. While the imprinting of *ZDBF2* is evident in humans and mice, whether this process is conserved in other mammals has not been addressed. Here, we show that the first exon of human *GPR1-AS* overlaps with that of a long terminal repeat (LTR) belonging to the MER21C subfamily of retrotransposons. Although this LTR family appears and is amplified in Boroeutherians, the magnorder of placental mammals that includes the Euarchontoglires and Laurasiatheria superorders, the MER21C insertion into the *GPR1-AS* orthologous region occurred specifically in the common ancestor of Euarchontoglires, a clade that includes extant primates, rodents, and rabbits. The first exon of mouse *Liz* does not overlap with an annotated LTR in standard repeat annotation; however, promoter activity assay and multiple sequence alignment suggests that it retains a functionally conserved relationship with the MER21C-overlapping first exon of *GPR1-AS*. Furthermore, directional RNA sequencing of placental tissues from rabbits and nonhuman primates also revealed *GPR1-AS* orthologs, with their first exon embedded within the same ancestral LTR. In contrast, allele-specific expression profiling of cow and tammar wallaby, mammals outside the Euarchontoglires group, revealed expression from both alleles in all tissues analyzed. Taken together, these observations suggest that imprinting of *ZDBF2* in Euarchontoglires had its genesis in the insertion of a MER21C element in their common ancestor. Our previous studies showed that LTRs reactivated in oocytes contribute to lineage-specific imprinting during mammalian evolution. The data presented here suggest that post-fertilization activation of an ancestral LTR-derived sequence can also contribute to the lineage-specific establishment of imprinted genes.

## INTRODUCTION

Genomic imprinting is a well-established biological phenomenon that refers to the differential expression of imprinted genes depending on their parental origin (Tucci et al. 2019; Kobayashi 2021). Epigenetic mechanisms, including DNA methylation, histone modification, and noncoding RNA molecules, regulate this process, establishing and maintaining parent-of-origin-specific gene expression patterns. Imprinted genes play vital roles in various biological processes such as fetal growth and development, placental function, metabolism, and behavior. These imprinted genes are often clustered within imprinted domains and regulated by epigenetic marks in specific imprinting control regions (ICRs). The inheritance of parent-of-origin-specific DNA methylation marks from the oocyte or sperm occurs in germline differentially methylated regions (germline DMRs), which are propagated in all somatic tissues of the next generation and function as ICRs (therefore, referred to as canonical imprinting). Imprinting patterns can vary significantly among species. Although some genes are imprinted in only one or a few species, others are conserved in many mammals. For example, nearly 200 imprinted genes have been identified in mice and humans. However, one comparative analysis suggested that fewer than half of these genes are imprinted in both species, suggesting the existence of many species- or lineage-specific imprinted genes (Tucci et al. 2019). The establishment of species-specific imprinting at specific loci can be driven by various evolutionary events, including retrotransposition of genes or transposable elements, which may lead to the acquisition of new genes as imprinted genes or the formation germline DMRs at CpG-rich regions (CpG islands)(Kobayashi 2021; Kaneko-Ishino and Ishino 2022). These CpG islands, often conserved as orthologous sequences across species, may become germline DMRs in the female germline as a consequence of transcription across the CpG island (Chotalia et al. 2009; Kobayashi et al. 2012a). Such transcription may be initiated within transposable elements which have integrated nearby. In such cases, the retrotransposition of genes or transposable elements is not directly responsible for the creation of CpG islands themselves but instead facilitates their recruitment into the imprinting system, contributing to the establishment of species-specific imprinting. Indeed, our previous studies have shown that oocyte-specific transcription from upstream promoters, such as integrated LTR retrotransposons, likely played a critical role in the establishment of lineage-specific maternal germline DMRs (Brind’Amour et al. 2018; Bogutz et al. 2019). An alternative form of imprinting involves the Polycomb marks H3K27me3 and H2AK119ub1, initially deposited in oocytes (Mei et al. 2021). Originally described in rodents, such “non-canonical” imprints are maternally inherited and in turn silence associated genes during preimplantation. Subsequently, these Polycomb marks are converted into DNA methylation over LTR elements, thus maintaining silencing of maternal alleles in the extraembryonic lineage (Hanna et al. 2019 ; Richard Albert et al. 2023). In addition, interspecies differences in the regulators of ICRs, such as *ZFP57* and *ZNF445*, as well as species-specific orthologs of DNA methyltransferases, including *DNMT3C,* are potential species-specific features: *ZFP57* and *ZNF445* cooperatively maintain ICR methylation, with *ZNF445* being more dominant in humans and *ZFP57* in mice; and *DNMT3C* is crucial for a paternally methylated DMR and appears to be specific to Muroidea. These features might also have contributed to the establishment of species-specific imprinted genes (Barau et al. 2016; Takahashi et al. 2019). However, these reports alone have not been sufficient to propose a comprehensive model for the emergence of all species-specific imprinted genes.

*Zinc finger DBF-type containing 2* (*ZDBF2*), a paternally expressed gene located on human chromosome 2q37.1, has been reported to regulate neonatal feeding behavior and growth in mice, although its molecular function remains unknown (Kobayashi et al. 2009; Glaser et al. 2022). There are at least three imprinted genes, *ZDBF2*, *GPR1* (alternatively named *CMKLR1*), and *ADAM23* around this locus (Hiura et al. 2010; Morcos et al. 2011). Recently, the *Zdbf2* gene was also shown to be paternally expressed in the rat, another member of the Muridae family (Richard Albert et al. 2023)(**Supplemental file 1**). Studies have shown that *ZDBF2* imprinting is regulated by two types of imprinted DMRs: a germline DMR located in the intron of the *GPR1* gene and a somatic (secondary) imprinted DMR upstream of *ZDBF2* (Kobayashi et al. 2012b; Kobayashi et al. 2013; Duffie et al. 2014). The germline DMR is methylated on the maternal allele, leading to silencing of *GPR1* antisense RNA (*GPR1-AS*, alternatively named *CMKLR1-AS*) expression on that allele. In contrast, the paternal allele is hypomethylated, allowing for the expression of *GPR1-AS*, which is transcribed in the reverse direction of *GPR1* (in the same direction as *ZDBF2*) from the germline DMR. Although it is not known whether *GPR1-AS* encodes a protein, a fusion transcript of *Gpr1-as* (*Platr12*) and *Zdbf2* (alternatively named *Zdbf2linc* or *Liz*; a long isoform of *Zdbf2*) has been identified in mice (Kobayashi et al. 2012b; Duffie et al. 2014). Notably, this fusion transcript exhibited an imprinted expression pattern similar to that of human *GPR1-AS* (**Supplemental file 1**). *Liz* transcription counteracts the H3K27me3-mediated repression of *Zdbf2* by promoting the deposition of antagonistic DNA methylation at the secondary DMR (Greenberg et al. 2017). Germline DMRs that function as ICRs generally maintain uniparental methylation throughout development; however, differential methylation of the germline DMR is no longer maintained at this locus after implantation because the paternally derived allele also becomes methylated after implantation. Thus, the secondary DMR, rather than germline DMR, is thought to maintain the imprinted status of this locus in somatic cell lineages. The unique regulatory mechanism responsible for imprinting at the *ZDBF2* locus appears to be conserved between humans and mice; however, whether this conservation extends to other mammals remains unknown.

Here, we focus on *GPR1-AS/Liz*, which is critical for the imprinting of *Zdbf2* in mice, and identified orthologous transcripts of *GPR1-AS* in primates and rabbits using RNA-seq-based transcript analysis. Strikingly, the first exon of these orthologs overlaps with a common LTR retrotransposon, suggesting that LTR insertion in a common ancestor led to the establishment of *ZDBF2* imprinting. In support of this hypothesis, *ZDBF2* expression is not imprinted in a mammalian outgroup to the Euarchontoglires that lack this LTR, including cattle and tammar wallabies. Taken together, our findings indicate that in the branch of the mammalian lineage that shows imprinting of *ZDBF2*, the key evolutionary event was the insertion at this locus of an LTR-derived sequence that becomes active only after fertilization.

## RESULTS

### Detection of *GPR1-AS* orthologs using RNA-seq data sets

*GPR1-AS* and *Liz* expression have been observed in placental tissues, blastocyst-to- gastrulation embryos, and/or ES cells in humans and mice (Kobayashi et al. 2009; Kobayashi et al. 2012b; Kobayashi et al. 2013; Duffie et al. 2014). Public human tissue RNA-seq data sets show that *GPR1-AS* transcription is detectable in the placenta, albeit at a relatively low level (RPKM value of around 0.45–1.8), but is significantly suppressed in other tissues (**Figure 1 – supplement figure 1A**) (Fagerberg et al. 2014; Duff et al. 2015). To determine whether *GPR1-AS* orthologs are present across different mammalian species, we performed transcript prediction analysis using short-read RNA-seq data from the placenta or specific extra-embryonic tissues of Eutheria (placental mammals) and Metatheria (marsupials).

**Figure 1.**
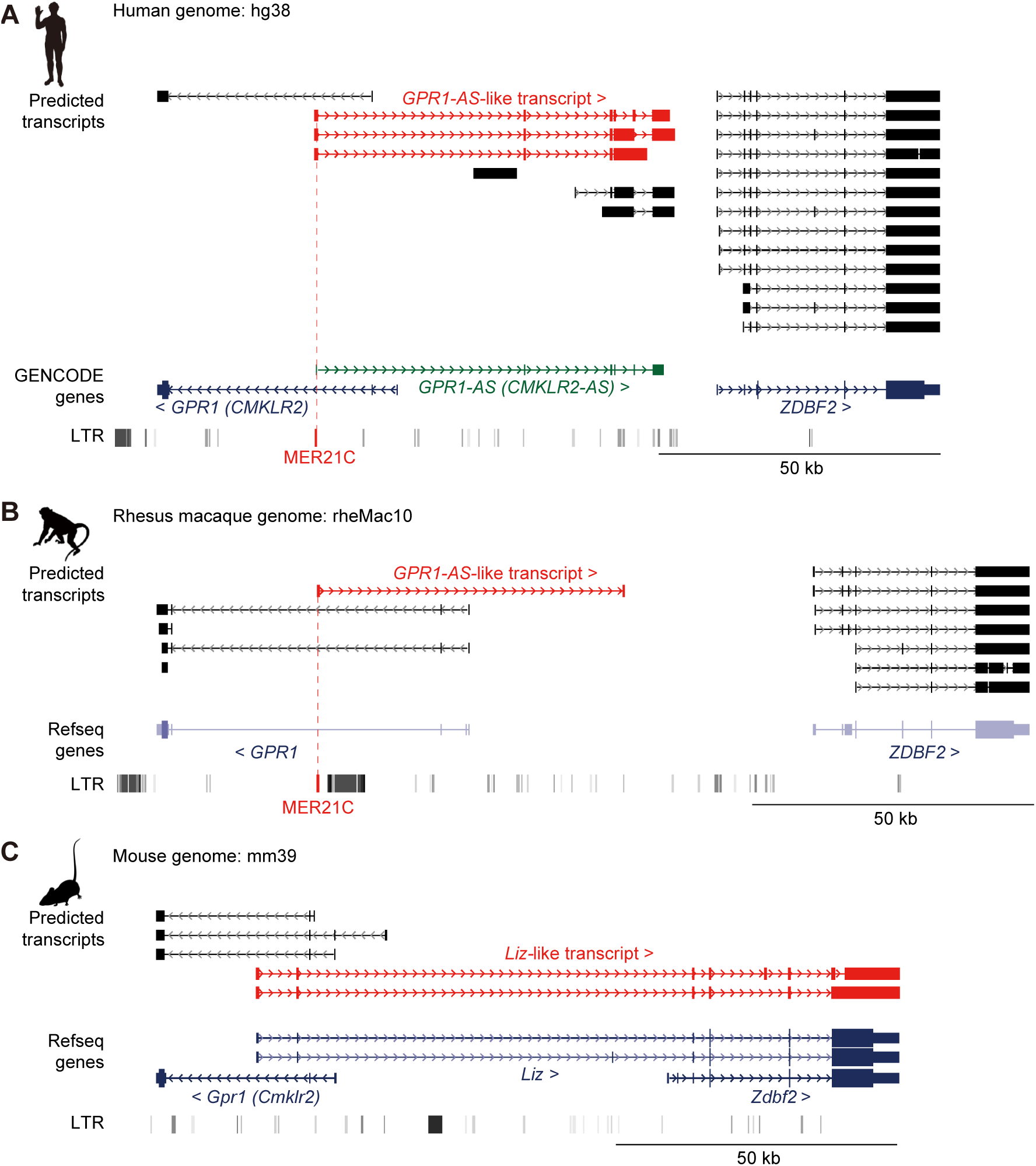
Identification of *GPR1-AS* orthologs from public placental transcriptomes. UCSC Genome Browser screenshots of the *GPR1*-*ZDBF2* locus in humans (**A**), rhesus macaques (**B**) and mice (**C**). Predicted transcripts were generated using public directional placental RNA-seq datasets (accession numbers: SRR12363247 for humans, SRR1236168 for rhesus macaques, and SRR943345 for mice) using the Hisat2-StringTie2 programs. Genes annotated from GENCODE or RefSeq databases and LTR retrotransposon positions from UCSC Genome Browser RepeatMasker tracks are also displayed. Among the gene lists, only the human reference genome includes an annotation for *GPR1-AS* (highlighted in green). *GPR1-AS*-like transcripts and MER21C retrotransposons are highlighted in red. Animal silhouettes were obtained from PhyloPic.

**Figure 2.**
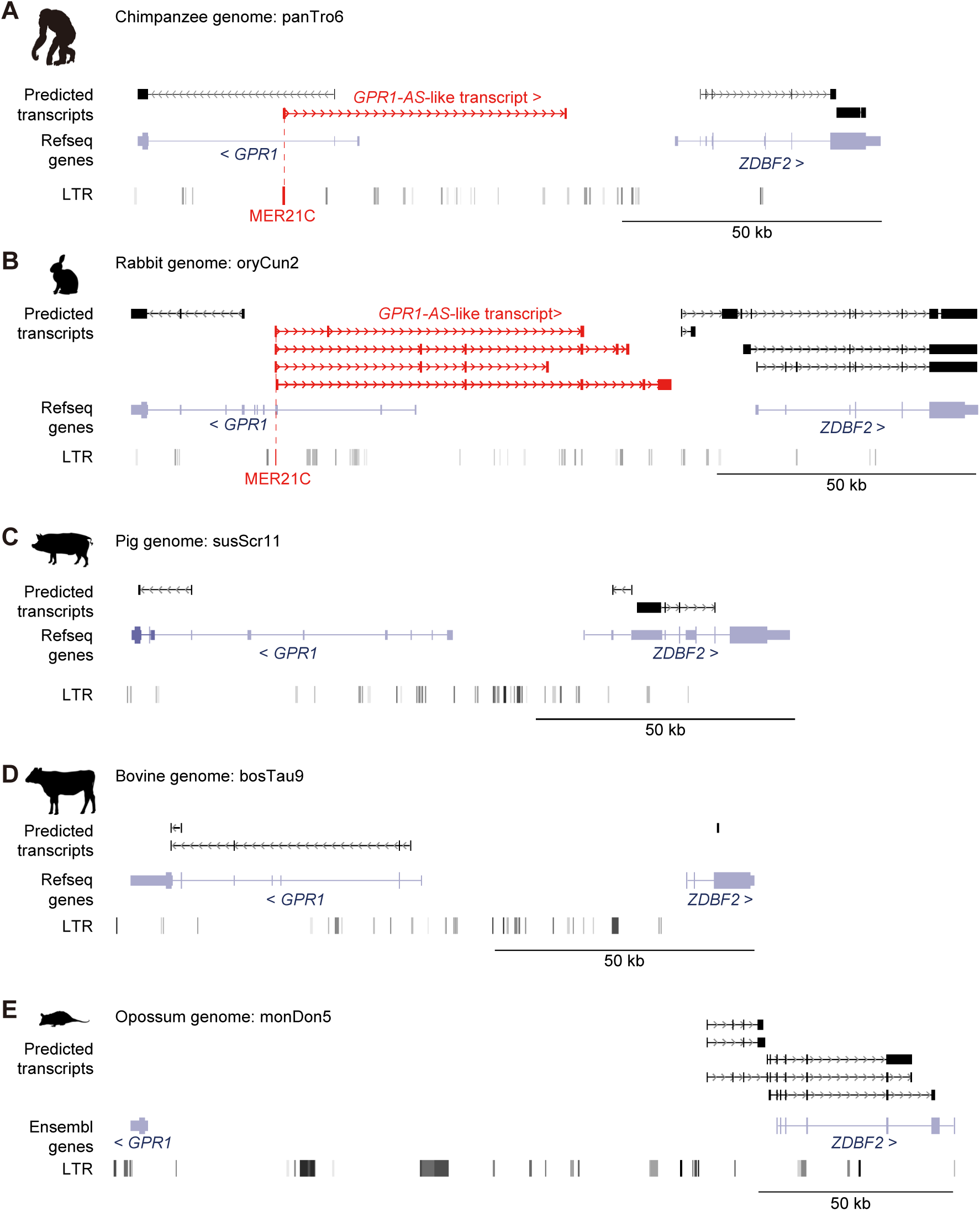
Identification of *GPR1-AS* orthologs from original placental and extra-embryonic transcriptomes. Predicted transcripts were generated from placental and extra-embryonic directional RNA-seq datasets of chimpanzee (**A**), rabbit (**B**), pig (**C**), cow (**D**), and opossum (**E**) with the Hisat2-StringTie2 programs. Genes annotated from RefSeq or Ensembl databases and their LTR positions are also shown. *GPR1-AS*-like transcripts and MER21C retrotransposons are highlighted in red.

First, we downloaded the placental RNA-seq data from 15 mammals: humans, bonobos, baboons, mice, golden hamsters, rabbits, pigs, cattle, sheep, horses, dogs, bats, elephants, armadillos, and opossums (Armstrong et al. 2017; Mika et al. 2022). These datasets were generated using conventional non-directional RNA-seq (also known as non-stranded RNA-seq), which does not retain information regarding the genomic element orientation of the sequenced transcripts. In addition, library preparation methods and the amount of data varied significantly among the datasets, ranging from millions to tens of millions (**Supplemental file 2**). Although transcriptional analysis of these datasets identified transcripts such as *GPR1-AS* in human and baboon placentas, the structures of these predicted transcripts were so fragmented that their full-length forms could not be robustly determined (**Figure 1 – supplement figure 1B-C**). Furthermore, we did not observe such a transcript at the *Gpr1*-*Zdbf2* locus in the mouse placental data, where *Liz* is known to be expressed. This suggests that conventional RNA-seq datasets may not be sufficient to accurately detect *GPR1-AS* in different species.

Next, we downloaded directional RNA-seq data (also known as strand-specific or stranded RNA-seq, which can identify the direction of transcription) from the human placenta, rhesus macaque trophoblast stem cells, and mouse embryonic placenta (Nesculea et al., 2014; Rosenkrantz et al. 2021). In each of these deeply sequenced datasets, which include a minimum of 50 million reads, transcripts originating from the *GPR1* intron were detected in the opposite direction to *GPR1*, extending towards the *ZDBF2* gene (i.e., *GPR1-AS* in humans and rhesus macaques, and *Liz* in mice) (**Figure 1**). Thus, *GPR1-AS* transcripts can be identified in placental tissues using deep directional RNA-seq data.

To identify *GPR1-AS*-like transcripts in other mammalian species, we generated deep directional RNA-seq datasets from the whole placenta of chimpanzees and the trophectoderm (TE) of rabbit, bovine, pig, and opossum post-gastrulation embryos, which yielded a total of 50–100 million reads (**Supplemental file 2**). Among these datasets, *GPR1-AS* orthologs were identified in chimpanzees and rabbits but were not observed in cows, pigs, or opossums (Figure 2). Additionally, we performed directional RNA-seq of tissues from the embryonic proper (embryonic disc: ED) of multiple individual animal embryos but did not identify any *GPR1-AS*-like transcripts in any of these samples (Figure 2 **– figure supplement 1**). Thus, rabbit *GPR1-AS* is expressed only in placental lineages. Identification of *GPR1-AS* in several primate species as well as mice and rabbits but not in cows, pigs, or opossums indicates that the regulatory region driving *GPR1-AS* transcripts likely originated in the common ancestor of the Euarchontoglires group.

### Origin of *ZDBF2* imprinting and *GPR1-AS* transcript

Since *GPR1-AS*/*Liz* is essential for *ZDBF2* imprinting in mice, it is assumed that *ZDBF2* imprinting system via *GPR1-AS* transcription is not established in other eutherians and marsupials that lack *GPR1-AS* (Greenberg et al. 2017). To test this hypothesis, we performed allele-specific expression analysis of the *ZDBF2* gene in embryonic, fetal and adult tissues of tammar wallabies and cattle, which are outside the Euarchontoglires group. We identified available SNPs at the 3’-UTR of *ZDBF2* in each of these mammals, and Sanger-sequencing-based polymorphism analysis showed the expression of both *ZDBF2* alleles in all bovine and wallaby tissues analyzed (**Figure 3A-B**)(**Supplemental file 1**). Similar analyses were performed using blood samples from rhesus macaques and rabbits, in which *GPR1-AS* was identified. Consistent with our findings in humans and mice, we observed paternal allele-specific expression of *ZDBF2* in these species (**Figure 3C-D**), in clear contrast to the bi-allelic expression observed in the embryonic, fetal, and adult tissues of tammar wallabies and cattle.

**Figure 3.**
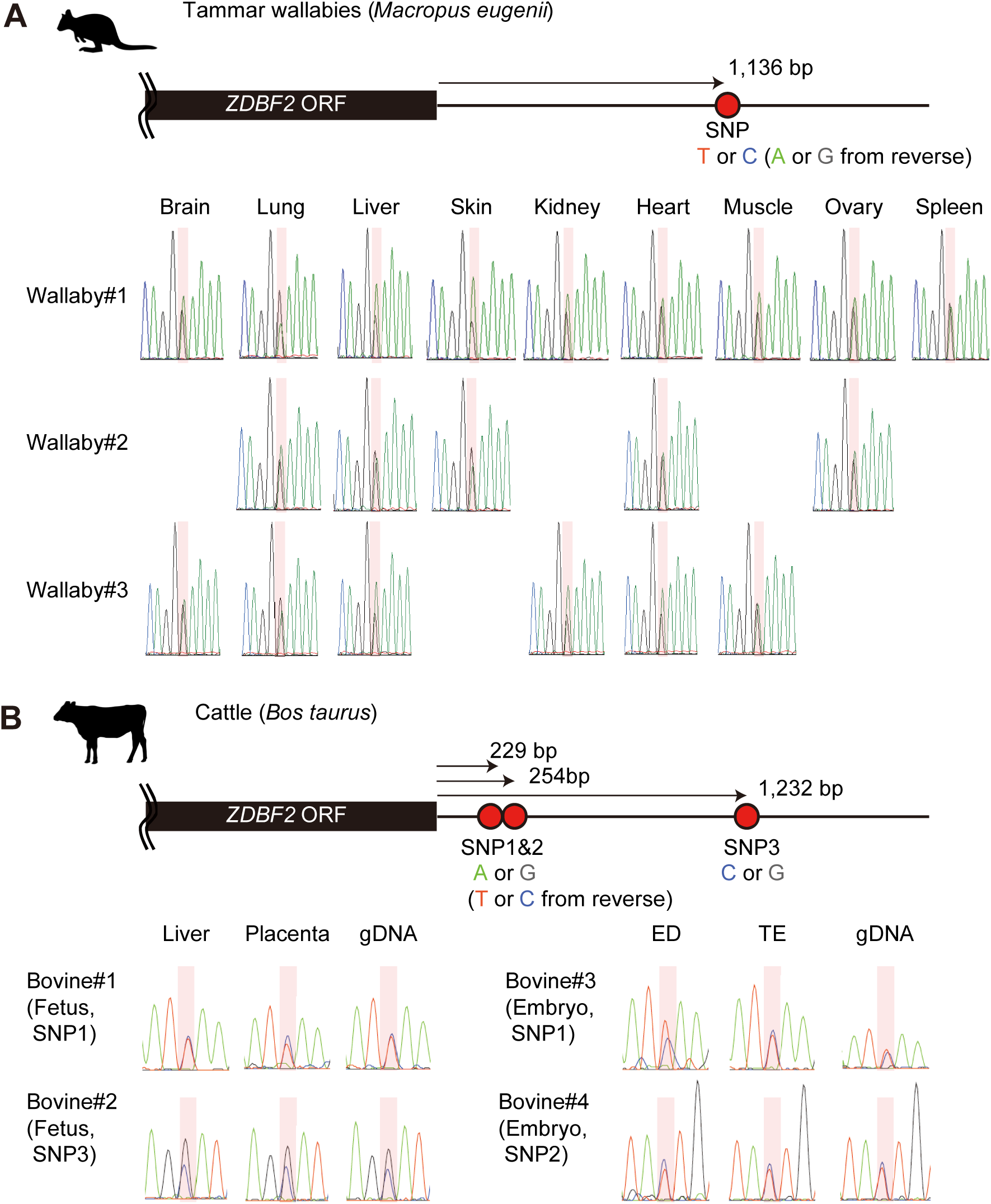
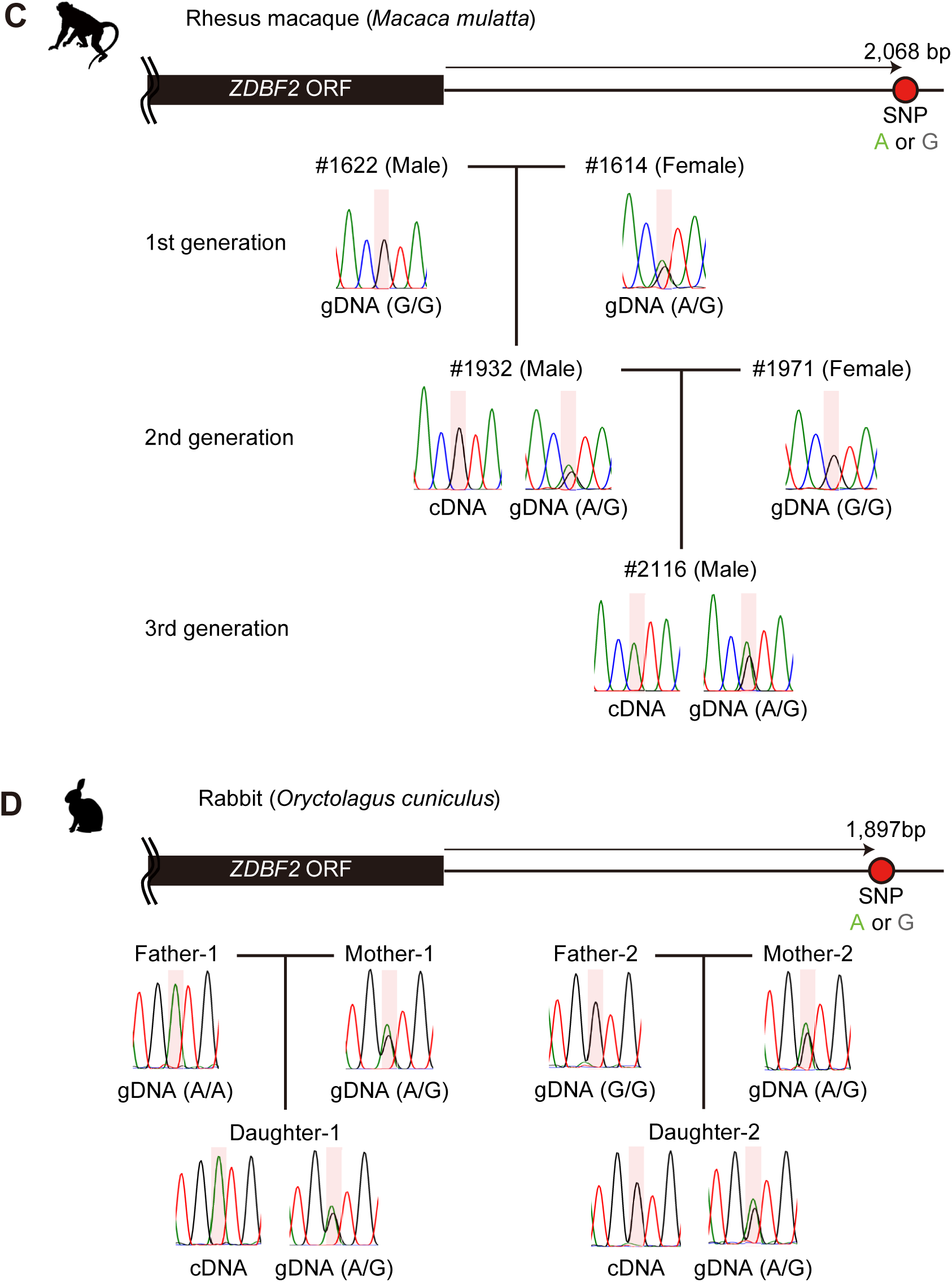
Allele-specific RT-PCR sequencing of *ZDBF2* in various mammals. Heterozygous genotypes were used to distinguish between parental alleles in adult tissues from tammar wallabies (**A**), fetal/embryonic tissues from cattle (**B**), blood samples from rhesus macaques (**C**) and rabbits (**D**), respectively. Primers were designed to amplify the 3’-UTR regions of *ZDBF2* orthologs and detect SNPs. Each SNP position is highlighted in red. Reverse primers were also used for Sanger sequencing.

To investigate a potential initiating mechanism in the germline for *ZDBF2* imprinting, which is established prior to the secondary DMR, we examined the presence of a germline DMR in both species with and without *ZDBF2* imprinting and *GPR1-AS* expression. Analysis of published whole-genome bisulfite sequencing data from rhesus macaque oocytes and sperm revealed DMRs between both germ cells, including an oocyte-methylated germline DMR at the first exon of *GPR1-AS* (**Figure 3 – figure supplement 3A**) (Gao et al. 2017). This finding aligns with previous observations in humans and mice (Kobayashi et al. 2012b; Kobayashi et al. 2013; Duffie et al. 2014). In contrast, analysis of published whole-genome bisulfite sequencing data for porcine and bovine oocytes and sperm (Ivanova et al. 2020) showed no oocyte-methylated germline DMRs in the *GPR1* intragenic region, where *GPR1-AS* is transcribed from an intron of *GPR1* in humans and rhesus macaques (**Figure 3 – figure supplement 3B**). Taken together, these results support the hypothesis that *ZDBF2* imprinting is restricted to mammals within the Euarchontoglires group.

Notably, the first exon of all *GPR1-AS* orthologs, except for mouse *Liz,* overlapped with an annotated MER21C element, a member of the LTR retrotransposon subfamily (**Figure 1, 2**, and **Figure 1 – figure supplement 1C**). While MER21C retrotransposons are broadly present in Boroeutherians, among the mammals tested, only primate and rabbit genomes have MER21C in the *GPR1* intron, where the first exon of *GPR1-AS* is located. Therefore, we compared the LTR sequence location information of the syntenic region of the *GPR1* and *GPR1-AS* loci to profile MER21C insertion sites in several mammalian genomes, excluding marsupials. Among eutherians, we found the insertion of MER21C (or MER21B retrotransposons that showed high similarity to MER21C) in the introns of *GPR1* in five groups of Euarchontoglires, including primates, colugos, treeshrews, rabbits, and rodents, with the exception of mouse, rat, and hamster (**Figure 4**). As the default parameters of RepeatMasker may not detect degenerate LTR sequences, we reanalyzed these three rodent species using less stringent RepeatMasker parameters but again failed to reveal the presence of an MER21C insertion in the relevant regions (**Figure 4 – figure supplement 1**). However, multiple genome alignments from UCSC/Cactus track indicated that the MER21C-derived sequence on the human *GPR1- AS* is likely conserved in mammals belong to Euarchontoglires, including mouse, rat, and hamster (**Figure 4 – figure supplement 2**) (Armstrong et al. 2020; Zoonomia 2020), albeit with a high level of degeneracy in the latter.

**Figure 4.**
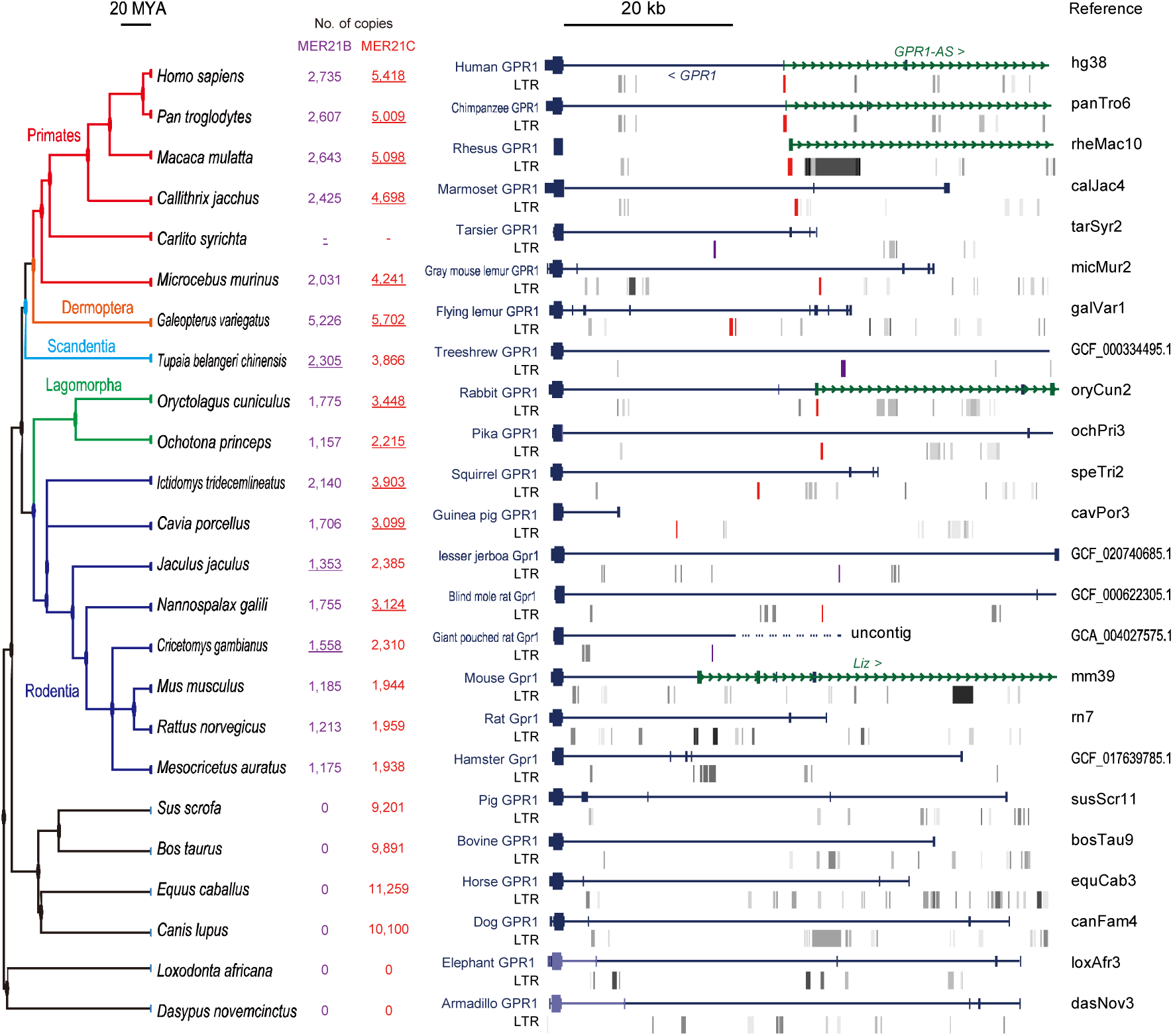
Multi-species comparison of LTR retrotransposon locations at *GPR1* locus. A total of 24 mammalian genomes were compared, including six primates (human, chimpanzee, rhesus macaque, marmoset, tarsier, and gray mouse lemur), one colugo (flying lemur), one treeshrew (Chinese treeshrew), two lagomorphs (rabbit and pika), eight rodents (squirrel, guinea pig, lesser jerboa, blind mole rat, giant pouched rat, mouse, rat, and golden hamster), and six other eutherians (pig, cow, horse, dog, elephant, and armadillo). Among the selected genomes, LTRs that can be considered homologous to MER21C, which corresponds to the first exon of *GPR1-AS*, are marked in red. In tarsier, treeshrew, lesser jerboa, and giant pouched rat, the orthologous LTRs were annotated as MER21B, which exhibits 88% similarity with MER21C in their consensus sequences through pairwise alignment. MER21B are marked in purple. According to Dfam, the MER21C and MER21B subfamilies are specific to the genomes of Boroeutherians and Euarchontoglires, respectively. The copy number of MER21C/B in selected species is shown in red and purple (LTRs likely matching the *GPR1-AS* exon are underlined). There are 5,418 and 2,529 copies of MER21C and 2,894 and 1,535 copies of MER21B in human and mouse genomes, respectively.

To analyze this syntenic region in greater detail, we compared MER21C sequences that overlapped with the first exon of *GPR1-AS* in each primate (345 bp, 339 bp, and 506 bp from humans, chimpanzees, and rhesus macaques, respectively) as well as rabbits (218 bp), with the first exon of *Liz* in mice (317 bp), and the common sequence of MER21C (938 bp). Pairwise alignment revealed high identities between primate sequences (identity: 71–72%) and the consensus MER21C sequence compared to rabbit (60%) and mouse sequences (46%) (**Figure 5 – figure supplement 1A**). The sequence identity in mice is so low that it cannot be distinguished from other LTRs or retrotransposons, which explains why it was not classified as MER21C by RepeatMasker analysis (**Figure 5 – figure supplement 1B-C**). Multiple sequence alignments and phylogenetic tree reconstruction reveal that the rabbit and mouse sequences deviate to a greater extent from the consensus MER21C sequence than the cluster formed by the primate sequences, indicating that a greater number of mutations have accumulated in the non-primate lineage of Euarchontoglires since their divergence from primates (**Figure 5A**). This observation aligns with the shorter generation time of lagomorphs and rodents compared to primates (Wu and Li 1985; Huttley et al. 2007).

**Figure 5.**
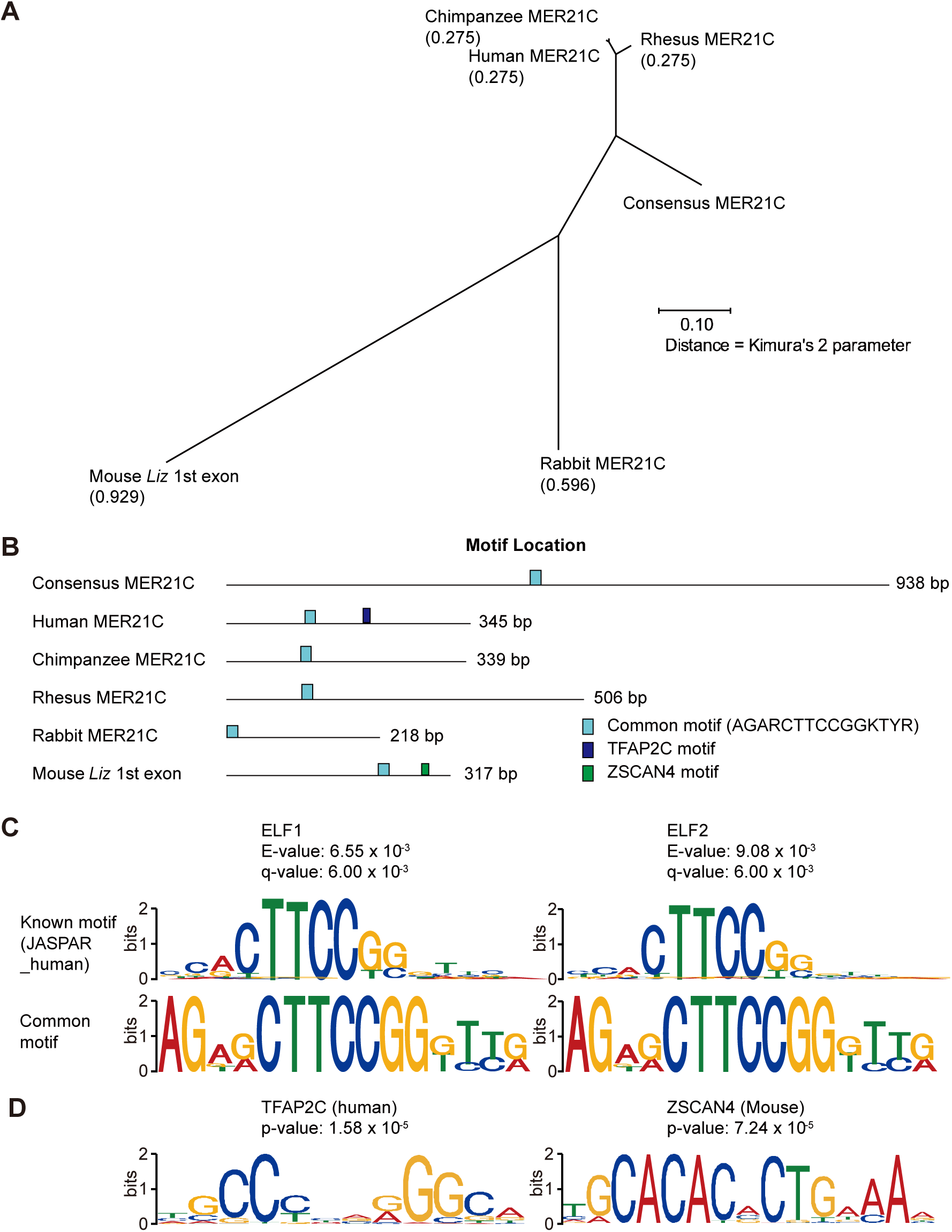
Comparison of MER21C-derived sequences overlapping the first exon of *GPR1- AS* orthologs. (**A**) Phylogenetic tree of MER21C-derived sequences estimated by multiple sequence alignment (MSA) using multiple sequence comparison by log-expectation (MUSCLE) program. (**B**) Positions of common and unique cis-acting elements at each sequence. (**C**) Motif structures of the common region that contains E74-like factor 1 and 2 (ELF1 and ELF2) binding motifs. (**D**) Motif structures of transcription factor AP-2 gamma (TFAP2C) and Zinc finger and SCAN domain containing 4 (ZSCAN4).

To evaluate whether the putative degenerate MER21C-derived regulatory region driving mouse *Liz* expression confers transcriptional activity in the mouse comparable to that of the MER21C element that drives human *GPR1-AS expression*, we performed a dual reporter assay in the human cell line HEK293T. Strikingly, the first exon of mouse *Liz* exhibits promoter activity greater than that of the human *GPR1-AS* promoter, despite the relatively low sequence similarity between the *Liz* first exon and the consensus MER21C sequence (**Figure 5 – figure supplement 2**). Taken together, these findings suggest that despite substantial sequence divergence, the functional elements responsible for initiating transcription of Liz in the mouse are derived from the same MER21C relic annotated in the human locus and can drive robust expression in human cells.

If this is indeed the case, then specific transcription factor binding motifs are likely shared between species in this regulatory region. To identify common cis-motifs, we compared these sequences with known transcription factor-binding motifs using the TOMTOM program in the MEME suite. Through this analysis, we found a common region that contained an ETS family transcription factor (ELF1 and ELF2) binding site, which had significant matches with E-values < 0.01 and q-values < 0.01 (**Figure 5B,C**). Additionally, we identified the TFAP2 and ZSCAN4 binding motifs (with p-values = 1.58 x 10^-5^ and 7.24 x 10^-7^) from the JASPAR database at the first exon of human *GPR1-AS* and mouse *Liz*, respectively (**Figure 5B,D**). Since *TFAP2C* and *ZSCAN4C* are activated in the placental lineage and preimplantation embryos, respectively (Zhang et al. 2019; Papuchova and Latos 2022) (**Figure 5 – figure supplement 3**), binding of these transcription factors may play a role in the transcriptional activation of *GPR1-AS* or *Liz* during embryogenesis, and in turn imprinting of *ZDBF2* .

### *GPR1-AS* transcription and LTR reactivation in human

*GPR1-AS* is reported to be expressed only briefly during embryogenesis, both before and after implantation, with the exception of the placental lineage. For instance, human *GPR1-AS* has been identified in ES cells, and mouse *Liz* expression has been observed in blastocyst-to-gastrulation embryos but not in gametes (Kobayashi et al. 2012b; Kobayashi et al. 2013; Duffie et al. 2014). Therefore, it is likely that the expression of *GPR1-AS*/*Liz* is activated during preimplantation. To investigate the timing of *GPR1-AS* activation during embryonic development, we obtained public human RNA-seq data from the oocyte to blastocyst stages and performed expression analysis and transcript prediction (Kai et al. 2022; Zou et al. 2022b). Among the datasets from oocytes; zygotes; 2-, 4-, and 8-cell embryos; inner cell mass (ICM); and TE from blastocysts, *GPR1-AS* was only detected at the 8-cell stage, ICM, and TE; *GPR-AS-like* transcripts were detected as predicted transcripts, with TPM values above 1 (**Figure 6**). This indicates that *GPR1-AS* expression begins at the 8-cell stage in humans. However, at the *GPR1*- *ZDBF2* locus, both *GPR1* and *ZDBF2* were expressed in oocytes, but only *GPR1* was downregulated following fertilization and became undetectable beyond the 8-cell stage (**Figure 6 and Figure 5 – figure supplement 3**). We also examined MER21C expression at each stage and found that it increased from the from 4-cell to 8-cell stage and peaked at the TE, coinciding with the appearance of *GPR1-AS* (**Figure 6 – figure supplement 1**). Moreover, focusing on Kruppel-associated box zinc-finger proteins (KRAB-ZFPs) involved in the suppression of transposable elements, we observed that *ZNF789*—a key KRAB-ZFP that binds MER21C—was specifically downregulated at the 4-cell stage (**Figure 5 – figure supplement 3**)(Imbeault et al. 2017). Taken together, these results suggest that *GPR1-AS* expression begins in the preimplantation embryo, concurrently with a transcription factor/repressor milieu permissive for expression from its MER21C-derived promoter.

**Figure 6.**
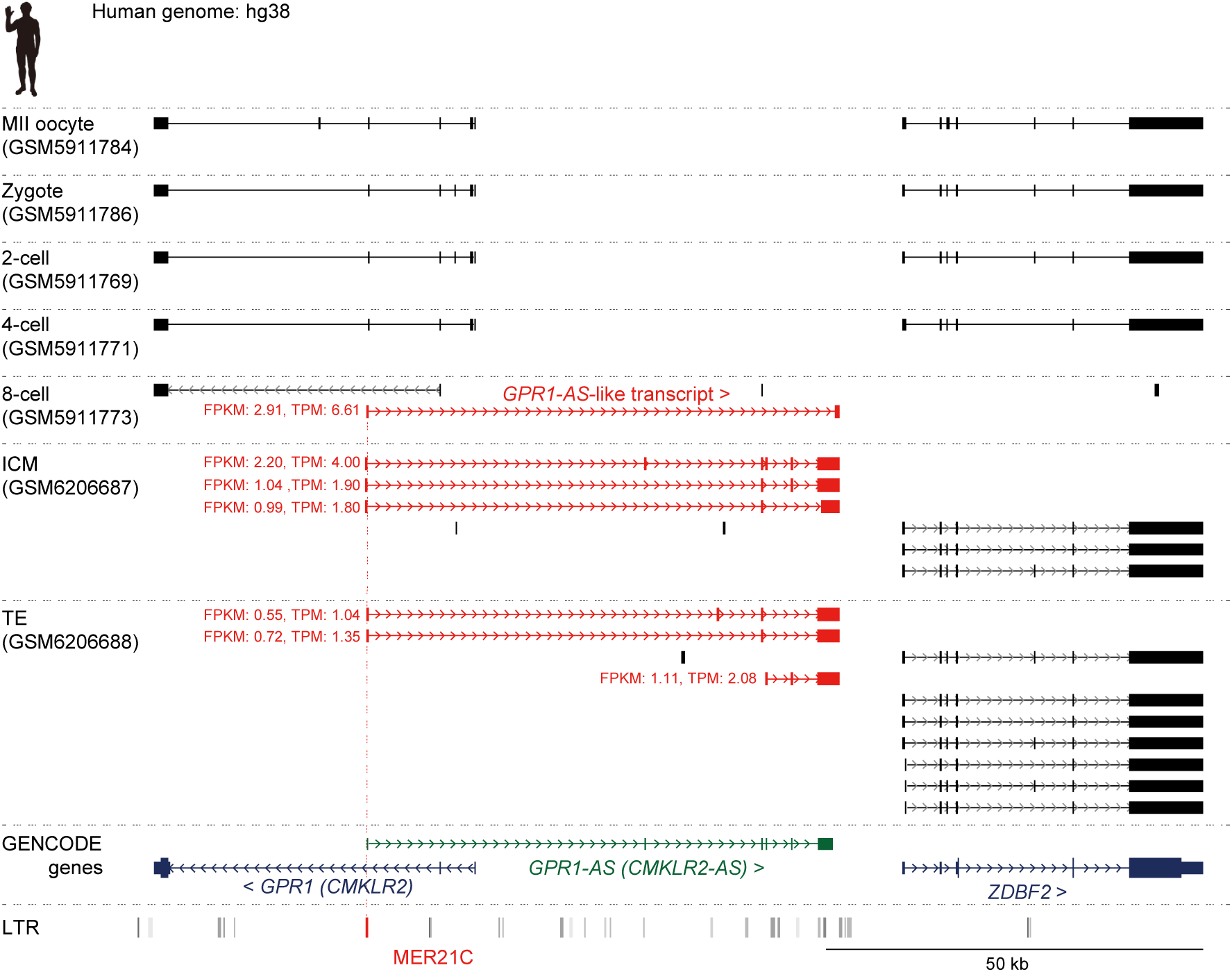
Initiation of *GPR1-AS* transcription before implantation. Genome browser screenshots of the *GPR1*-*ZDBF2* locus in humans at preimplantation stages, including the MII oocyte, zygote, 2-cell, 4-cell, 8-cell, inner cell mass (ICM), and trophectoderm (TE) from the blastocyst. Predicted transcripts were generated from publicly available full-length RNA-seq datasets, with detected *GPR1-AS*-like transcripts and their FPKM and TPM values highlighted in red.

To determine whether specific epigenetic marks are associated with the LTR-derived promoter located in the first exon of human *GPR1-AS* and mouse *Liz*, we surveyed the ChIP-Atlas database (Zou et al. 2022a) (**Figure 7 – figure supplement 1**). Both sequences that constitute the oocyte-derived germline DMR exhibit an enrichment of H3K4me3 in sperm or male germ cells, consistent with DNA hypomethylation in sperm. These sequences also show an enrichment of H3K27ac in pluripotent stem cells (such as ES and iPS cells). Although as discussed above, the first exon of mouse *Liz* has not been computationally determined as an LTR (according to the RepBase database), the mouse sequence is also enriched for H3K9me3, a repressive mark which is deposited at retrotransposons, in multiple cell types and tissues. Additionally, human and mouse germline DMRs coincided with the TFAP2C peak (trophoblast stem cells) and ZSCAN4C peak (ES cells), respectively, and aligned with the binding motif predictions found in the JASPAR database (**Figure 5B,D**). These data indicate that the first exons of human *GPR1-AS* and mouse *Liz* exhibit similar epigenomic modification variations from gamete to embryonic development, and their promoter activities are triggered after fertilization. However, the specific regulatory pathways governing this activation may differ between species, likely reflecting differences in transcription factor involvement and regulatory networks (**Figure 7**).

**Figure 7.**
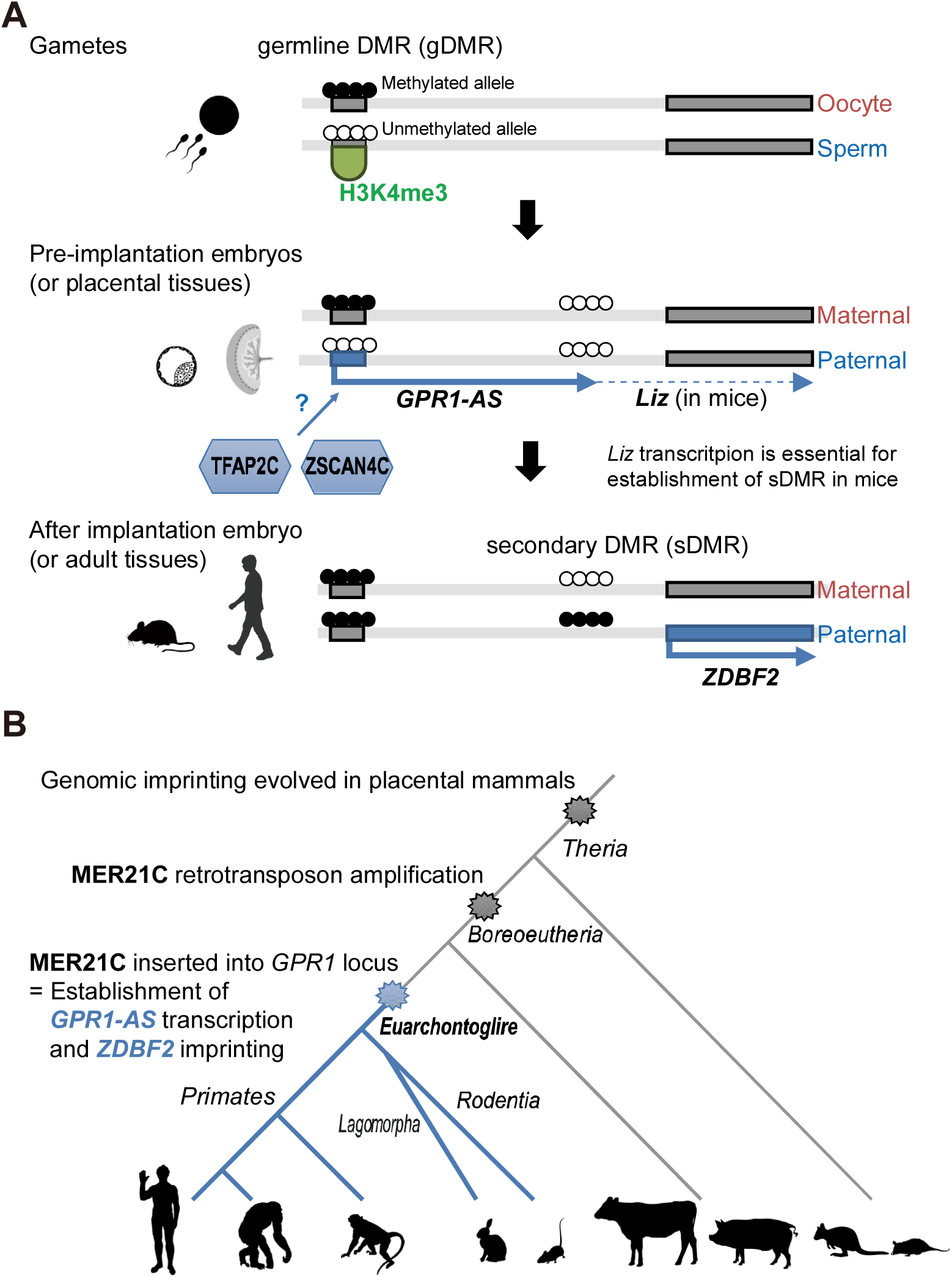
Establishment of *ZDBF2* imprinted domain in evolution and genome biology. (A) Scheme of epigenetic and transcriptional changes at the first exon of mouse *Liz* and human *GPR1-AS*. (B) Timescale of the evolution of *ZDBF2* imprinting and LTR (MER21C) insertion.

## DISCUSSION

The *ZDBF2* gene, the last canonical imprinting gene identified in mice and humans, was recently shown to regulate neonatal feeding behavior (Glaser et al. 2022). This discovery has garnered attention owing to its implications for behavioral phenotypes, aligning with the conflicting hypothesis of genomic imprinting mechanisms. The conflict hypothesis proposes that the evolutionary significance of genomic imprinting mechanisms lies in the functional conflicts between paternally and maternally expressed genes, resulting from the conflicting genomic survival strategies of the father, child, and mother (Moore and Haig 1991). *ZDBF2* does not directly influence cell differentiation or ontogeny, such as embryogenesis, but rather affects individual growth through an indirect pathway involving feeding behavior. In humans, “imprinting diseases”, such as Prader-Willi syndrome and Angelman syndrome, cause psychiatric disorders and growth retardation (Nicholls 2000). Considering that many imprinted genes show specialized expression patterns in the brain, *ZDBF2* is particularly interesting as a factor that can induce behavioral effects. However, the evolutionary origin of *ZDBF2* imprinting had remained unclear. In this study, we report that *ZDBF2* does not exhibit imprinted expression in cattle and tammar wallabies, mammalian species that do not belong to the Euarchontoglires clade. In contrast, we identified *GPR1-AS* orthologs in all the Euarchontoglires species examined, including humans, chimpanzees, baboons, rhesus macaques and rabbits. We propose that the first exon of *GPR1-AS* is derived from the MER21C retrotransposon and that the insertion of this LTR at the *GPR1* intragenic region in the common ancestor of Euarchontoglires was the crucial event in the genesis of imprinting of the *ZDBF2* gene (**Figure 7**).

Two paternally expressed imprinted genes, *PEG10*/*SIRH1*, and *PEG11*/*RTL1*/*SIRH2*, that encode GAG-POL proteins of *sushi-ichi* LTR retrotransposons have been identified in mammals and are essential for placenta formation and maintenance (Kaneko-Ishino and Ishino 2012). The presence of these genes provides evidence that LTR insertions have played a significant role in genomic imprinting as well as placentation throughout evolution of placental mammals. In contrast, while the *ZDBF2* gene is present in all vertebrate genomes, it is only imprinted in mammals belonging to the Euarchontoglires superorder of placental mammals. This suggests that a nonimprinted gene acquired imprinting status through the insertion of an LTR retrotransposon. Indeed, we previously demonstrated that transcriptional activity of specific LTRs in oocytes results in species-specific imprinting of numerous genes in rodents and primates (Bogutz et al. 2019). These LTRs act as alternative promoters and produce unique transcripts in the oocyte, which we referred to as LITs (LTR-initiated transcripts). Such transcription leads to a high level of methylation in the transcribed region, similar to gene body methylation, and is responsible for extensive interspecies differences in DNA methylation (Brind’Amour et al. 2018). The key difference between these LTRs and the LTR retrotransposon focused on here is the stage in development when LTR (re)activation occurs, i.e. before or after fertilization, respectively. Some LITs, which are driven by LTR families that are particularly active in oocytes, become active during oogenesis and establish oocyte-derived gDMRs of species-specific imprinted genes, such as *Impact* and *Slc38a4* genes in mice. Strikingly, deletion of these LTRs causes loss of imprinting in mouse (rodent)- specific imprinted genes (Bogutz et al. 2019). *GPR1-AS* is a (likely noncoding) RNA whose transcription is initiated from an LTR-derived sequence, similar to the LITs found in oocytes; however, transcription in this case starts after fertilization. In mice, *Liz*, a long isoform of *Zdbf2* and putative *Gpr1-as* ortholog, is essential for establishing a somatic secondary DMR upstream of *Zdbf2* (Greenberg et al. 2017). Although the first exon of *Liz* does not overlap with an annotated LTR, MER21C (or MER21B, a closely related LTR family) insertions are found in the homologous region in most rodents, with the exception of mice, rats, and hamsters. Our analyses suggest that a MER21C relic is also present in these rodents, but has accumulated a number of mutations and in turn is no longer recognized as a MER21C element using optimal pairwise alignment algorithms such as RepeatMasker. Indeed, multiple genome alignment using Cactus reveals that the first exon of human *GPR1-AS* is homologous across all selected rodents, including mouse, rat, and hamster. Based on these observations, we propose that *Liz* and *GPR1- AS* are bona fide LITs with a MER21C-derived sequence serving as an alternative promoter. A combined approach involving the annotation of repetitive elements with tools like RepeatMasker and the reconstruction of ancestral genomes using multiple-genome alignments can uncover highly degenerated LTR relics. For instance, Didier Trono’s group recently identified numerous previously unannotated transposable elements in the human genome using a similar methodology (Matsushima et al. 2024). Advances in such computational analyses could enable the identification of additional functional LTR relics and provide insights into the origin of lineage-specific regulatory sequences that are embedded within newly discovered lineage-specific LTR relics.

Transposable elements are recognized by DNA-binding factors and their co-repressors and are suppressed through epigenetic and post-transcriptional regulation in both germline and somatic tissues to protect host genome stability. For example, LTR-retrotransposons are bound by KRAB-ZFPs, which recruit the KAP1/SETDB1 co- repressor complex, promoting deposition of the repressive histone modification H3K9me3 (Matsui et al. 2010; Karimi et al. 2011). However, numerous retrotransposons are activated and robustly expressed during maternal to zygotic transition, where dynamic epigenetic reprogramming occurs. Indeed, a subset of transposable elements have been co-opted by the host during this window of development, including in support of embryonic development (Sakashita et al. 2023). In humans, retrotransposons are activated at the 8-cell stage and gradually downregulated in later developmental stages, coinciding with embryonic genome activation. In addition to the full-length endogenous retrovirus (ERV) sequences with partial coding potential, approximately 85% of human ERVs (HERVs) exist as solitary LTRs, known as “solo LTRs,” which provide a rich source of cis-regulatory elements for gene expression during human embryonic development. Retrotransposons located near host genes can act as alternative promoters or regulatory modules, such as enhancers, to activate embryonic genes (Grow et al. 2015; Hashimoto et al. 2021). The discovery of the *GPR1-AS* as a transcript that initiates in an ancestral LTR reveals a new aspect of the functional role of these ectopic regulatory elements, with activation in this case occurring after fertilization.

In summary, this study reveals that the origin of imprinting of *ZDBF2* can be traced back to a common ancestor of Euarchontoglires, in which an LTR element inserted in the locus. The solo LTR derived from this element (following recombination between 5’ and 3’ LTRs) drives expression of *GPR1-AS*, which in turn plays a critical role in establishing imprinting of the *ZDBF2* locus, including in both mice and humans. Although the house mouse is used widely in the laboratory, and the *Mus musculus* genome has been extensively characterized, the retroviral origin of this promoter would not have been identified if this analysis had been performed solely on mice, as the annotated exon 1 of *Liz* (alternatively named *Gpr1-as, Platr12*, or *Zdbf2linc*) is highly degenerate and in turn not recognized as an LTR in this species. Comparative analysis of multi-omics data, including genomes, transcriptomes, and epigenomes, across species was critical for the identification of the central role of a MER21C LTR in imprinting of *ZDBF2*. This study adds to a growing list of LTR elements that have been domesticated in mammals, with their transcriptional activity serving as a mechanism for the genesis of imprinting, including for a number of genes during gametogenesis and in the case of *ZDBF2,* following fertilization.

## METHODS

### Ethical approval for animal work

Animal experiments (chimpanzee, rhesus macaque, rabbit, pig, cow, and opossum) were conducted in accordance with the guidelines of the Science Council of Japan and approved by the Institutional Animal Care and Use Committee of the University of Tokyo, the National Institute for Physiological Sciences, RIKEN Kobe Branch, Meiji University, Shinshu University, Hokkaido University, the Center for the Evolutionary Origins of Human Behavior of Kyoto University, and Nara Medical University. Experimental procedures of tammar wallaby conformed to Australian National Health and Medical Research Council (2013) guidelines and were approved by the Animal Experimentation Ethics Committees of the University of Melbourne.

### Animals for transcriptome analysis

Placental samples from chimpanzees (*Pan troglodytes verus*) were provided by Kumamoto Zoo and Kumamoto Sanctuary via the Great Ape Information Network. The samples were stored at −80 °C before use. Total RNA was isolated from the placenta using an AllPrep DNA/RNA Mini Kit (Qiagen, Netherland). Bovine (*Bos taurus*, Holstein cow) embryos were prepared by in vitro oocyte maturation, fertilization (IVF), and subsequent in vitro embryo culture (Akizawa et al. 2018). Briefly, presumptive IVF zygotes were denuded by pipetting after 12 h of incubation and cultured up to the blastocyst stage in mSOFai medium at 38.5 °C in a humidified atmosphere of 5% CO_2_ and 5% O_2_ in air for 8 days (D8). D8 blastocysts were further cultured on agarose gel for 4 dayss as described previously (Akizawa et al. 2018; Saito et al. 2022). The embryo proper (embryonic disc: ED) and TE portions of IVF D12 embryos were mechanically divided using a microfeather blade (Feather Safety Razor, Japan) under a stereomicroscope. Total RNA from ED and TE samples was isolated using the ReliaPrepTM RNA Cell Miniprep System (Promega, MA, USA) according to the manufacturer’s instructions. Rabbit embryos were prepared on embryonic day 6.75 (E6.75) by mating female Dutch and male JW rabbits (Kitayama Labes, Japan). Pig embryos (*Sus scrofa domesticus*: Landrace pig) at E15 and opossum embryos at E11 were prepared (Kiyonari et al. 2021). Individual embryo proper and TE were mechanically divided using a 27-30G needle and tweezer under a stereomicroscope with 10% FCS/M199 -medium in rabbits, HBSS medium in pigs, and 3% FCS/PBS in opossums. Total RNA was isolated from individual tissues using the RNeasy Mini and Micro Kit (Qiagen). The quality of the total RNA samples was assessed using an Agilent 2100 Bioanalyzer system (Thermo Fisher Scientific, MA, USA) and high-quality RNA samples (RIN≧7) were selected for RNA-seq library construction.

### Strand-specific RNA library preparation and sequencing

Embryonic total RNA (10 ng) from opossums and pigs, 5 ng of placental total RNA from chimpanzees, and 1 ng of embryonic total RNA from rabbits and cattle were reverse transcribed using the SMARTer Stranded Total RNA-Seq Kit v2 - Pico Input Mammalian (Takara Bio, Japan) according to the manufacturer’s protocols, where Read 2 corresponds to the sense strand due to template-switching reactions. RNA-seq libraries were quantified by qPCR using the KAPA Library Quantification Kit (Nippon Genetics, Japan). All libraries were pooled and subjected to paired-end 75 bp sequencing (paired-end 76 nt reads with the first 1 nt of Read 1 and the last 1 nt of Read 2 trimmed) using the NextSeq500 system (Illumina, CA, USA). For each library, the reads were trimmed using Trimmomatic to remove two nucleotides from the 5’ end of the Read 2.

### RNA-seq data set download

Strand-specific RNA-seq datasets from the human bulk placenta, rhesus macaque trophoblast stem cells, and mouse bulk placenta at E16.5, were downloaded from the accession numbers SRR12363247 and SRR12363248 for humans, SRR1236168 and SRR1236169 for rhesus macaques, and SRR943345 for mice (Necsulea et al. 2014; Rosenkrantz et al. 2021). Non-directional RNA-seq data from placentas or extra- embryonic tissues of 15 mammalian species, including humans, bonobos, baboons, mice, golden hamsters, rabbits, pigs, cattle, sheep, horses, dogs, bast, elephants, armadillos, and opossums, were downloaded (Armstrong et al. 2017; Mika et al. 2022).

Full-length RNA-seq (based on the smart-seq method) data from human oocytes, zygotes, 2-, 4-, and 8-cell, ICM, and TE were downloaded from previous studies (Kai et al. 2022; Zou et al. 2022b). The corresponding accession numbers are shown in **Supplemental file 2**.

### RNA-seq data processing

All FASTQ files were quality-filtered using Trimmomatic and mapped to individual genome references (**Supplemental file 2**) using Hisat2 with default parameters and Stringtie2 with default parameters and “-g 500” option. Fragments per kilobase million (FPKM) and transcripts per kilobase million (TPM) were calculated using StringTie2 with “-e” option and the GENCODE GTF file, selecting lines containing the "Ensembl_canonical" tag. Read counts of human transposable element were computed at the subfamily level using the TEcount. The read counts were normalized by the total number of mapped reads to retrotransposons in each sample as Reads per kilobase million (RPKM).

### Data visualization

All GTF files (StringTie2 assembled transcripts) were uploaded to the UCSC Genome Browser, and the predicted transcripts, annotated genes (GENCODE, Ensembl, or RefSeq), and LTR locations were visualized. The expression profiles of human *GPR1*, *GPR1-AS*, and *ZDBF2* in the somatic tissues (Fagerberg et al. 2014; Duff et al. 2015), and these imprinted genes, the associated transcription factors (*TFAP2C*, *ZSCAN4*, *ELF1*, and *ELF2*), and LTR subfamilies in early embryo (Kai et al. 2022; Zou et al. 2022b) were downloaded or calculated, and visualized as heatmaps using the Morpheus software. The estimated evolutionary distance between the selected mammals was visualized as a physiological tree with a branch length of millions of years (MYA) using Timetree. The pairwise alignment, multiple sequence alignment, and physiological tree of MER21C sequences overlapping *GPR1-AS* were conducted with Genetyx software (Genetyx, Japan) using the MSA and MUSCLE programs. The common cis-motif was identified and compared with known transcription factor binding motifs from JASPAR CORE database using the XSTREAM and TOMTOM programs in the MEME suite. JASPAR motifs, logos, and *p*-values were obtained from the JASPAR hub tracks of the UCSC Genome Browser and JASPAR database. LTR positions of human (hg19) and mouse (mm10) were downloaded from UCSC Genome Browser and re-uploaded to Integrative Genomics Viewer with epigenetic patterns from ChIP-Atlas (Zou et al. 2022a) and oocyte/sperm DNA methylomes (Brind’Amour et al. 2018).

### Allele-specific expression analysis

Tammar wallabies (*Macropus eugenii*) of Kangaroo Island origin were maintained in our breeding colony in grassy, outdoor enclosures. Lucerne cubes, grass and water were provided ad libitum and supplemented with fresh vegetables. Pouch young were dissected to obtain a range of tissues. DNA/RNA was extracted from multiple tissues of the Tammar wallaby using TRIzol Reagent (Thermo Fisher Scientific), and cDNA synthesis was performed using Transcriptor First Strand cDNA Synthesis Kit (Roche, Switzerland). PCR was performed using TaKaRa Ex Taq Hot Start Version (TaKaRa Bio) with primers designed at 3’UTR of wallaby *ZDBF2* according to the manufacturer’s instructions. Fetal tissues of pregnant healthy Holstein cattle were obtained from a local slaughterhouse. DNA/RNA was extracted from bovine fetus (liver), placentas, and *in vitro* cultured embryos using ISOGEN (Wako, Japan), and cDNA synthesis and PCR were performed using SuperScript IV One-Step RT-PCR System (Thermo Fisher Scientific) with primers designed at 3’UTR of bovine *ZDBF2* according to the manufacturer’s instructions. Target regions were also amplified using genomic DNA and Takara EX Taq Hot Start Version (Takara Bio). Whole blood samples of rhesus macaque (*Macaca mulatta*) and rabbit (Japanese White, *Oryctolagus cuniculus*) were obtained from PRI of Kyoto University and purchased from Oriental Bio Service (Japan), respectively. DNA/RNA was extracted from blood samples using ISOSPIN Blood & Plasma DNA (Nippon Gene, Japan) and Nucleospin RNA Blood (Takara Bio) and cDNA synthesis and PCR were performed using SuperScript IV One-Step RT-PCR System (Thermo Fisher Scientific) with primers designed at 3’UTR of rhesus macaque and rabbit *ZDBF2* according to the manufacturer’s instructions. Sanger sequencing was performed using a conventional Sanger sequencing service (Fasmac, Japan) and ABI 3130xl Genetic Analyzer (Thermo Fisher Scientific). Sanger sequencing results were visualized using 4Peaks. Primers used in this analysis are listed in **Supplemental file 3**.

### DNA Methylome Analysis

The DSS R package (v2.54.0) was used to identify DMRs from oocyte and sperm CpG report files (accession numbers GSM1466810, GSM1466811, and GSE143849), with smoothing enabled and customized parameters. These parameters included a minimum CpG methylation difference of 50%, a p-value threshold of 0.05, a minimum DMR length of 200 bp, and other default settings. All BED files were uploaded to the UCSC Genome Browser and CpG sites with DNAme levels and called DMR positions were visualized.

### Dual reporter assay

Luciferase reporter plasmids were constructed using the pGL4.13 vector (Promega). Sequences containing the first exon and upstream regions of both mouse *Liz* and human *GPR1-AS* were amplified by PCR using TaKaRa HS Perfect Mix (Takara Bio) and specific primers with NheI and HindIII recognition sites at their 5’ ends (**Supplemental file 3**). The PCR products were inserted between the NheI and HindIII sites of the pGL4.13 vector using DNA Ligation Mix (Takara Bio). To evaluate the promoter activity of these sequences, the SpectraMax DuoLuc Reporter Assay Kit (Molecular Devices, CA, USA) was employed. Human embryonic kidney 293T (HEK293T) cells were cultured in Dulbecco’s Modified Eagle’s Medium (Thermo Fisher Scientific, Cat. No. 10313021) supplemented with 10% fetal bovine serum (NICHIREI Biosciences, Tokyo, Japan), 0.1 mg/mL penicillin-streptomycin (Thermo Fisher Scientific), and GlutaMax (Thermo Fisher Scientific, Cat. No. 35050061), on gelatin-coated dishes. For luciferase reporter assays, HEK293T cells were seeded in gelatin-coated 96-well plates at a density of 4 × 10⁵ cells/well prior to transfection. Transient transfections were carried out using Lipofectamine 2000 Transfection Reagent (Thermo Fisher Scientific) according to the manufacturer’s instructions. Each well was co-transfected with 10 ng of the pGL4.75 plasmid (Promega) as an internal transfection control reporter and 100 ng of the pGL4.13 plasmid with or without the candidate regulatory elements. After 48 hours of transfection, dual-luciferase assays were performed as per the manufacturer’s protocol (Molecular Devices). Luminescence signals were measured using a SpectraMax iD3 Multi-Mode Microplate Reader (Molecular Devices). Firefly luciferase activities (pGL4.13) were normalized to Renilla luciferase activities (PGL4.75).

### Softwares used

Trimmomatic (http://www.usadellab.org/cms/?page=trimmomatic)

Hisat2 (http://daehwankimlab.github.io/hisat2/)

StringTie2 (https://ccb.jhu.edu/software/stringtie/)

TEcount (https://github.com/bodegalab/tecount/)

Integrative Genomics Viewer (IGV: https://software.broadinstitute.org/software/igv/)

4Peaks (https://nucleobytes.com/4peaks/)

DSS (https://bioconductor.org/packages/release/bioc/html/DSS.html)

### Web-tools used

UCSC Genome Browser (https://genome.ucsc.edu/)

Morpheus (https://software.broadinstitute.org/morpheus/)

Timetree (http://timetree.org/)

MEME Suite (https://meme-suite.org/meme/)

JASPAR (https://jaspar.genereg.net/)

ChIP-Atlas (https://chip-atlas.org/)

PhyloPic (https://www.phylopic.org/)

RepeatMasker (https://www.repeatmasker.org/)

PipMaker and MultiPipMaker (http://pipmaker.bx.psu.edu/pipmaker/)

## DATA ACCESS

All raw sequencing data generated in this study were submitted as FASTQ files to the NCBI SRA (https://www.ncbi.nlm.nih.gov/sra). The accession numbers are shown in **Supplemental file 2**.

## ACKNOWLEDGEMENTS

H. K. and K. K. were supported by MEXT KAKENHI, Grant Numbers JP21H02382 and JP20H00471, respectively. H.K. and H.I. were supported by The Cooperative Research Programs of the NODAI Genome Research Center at Tokyo University of Agriculture, and the Center for the Evolutionary Origins of Human Behavior and Wildlife Research Center at Kyoto University. H.K. received additional support from the Mitsubishi Foundation and the Takeda Science Foundation. Additionally, we acknowledge Mr. Yasuyuki Osada (Kitayama Laboratories, Japan) for his technical assistance in handling the rabbit embryos, and Mr. Aaron Bogutz (University of British Columbia) for his helpful comments.

## Author Contribution

Designed the study and wrote the manuscript: H. Kobayashi

Data analysis: H. Kobayashi., T.I.

NGS data processing: S.K., K.T., T.T

Dual reporter assay: H. Kobayashi, S.N.

Wallaby sample preparation and data analysis: S. Suzuki, M.H., M.B.R.

Bovine sample preparation: S. Saito, M. K.

Rabbit sample preparation: T.K.

Pig sample preparation: H.N., H.M., K. Nakano, A.U.

Opossum sample preparation: H. Kiyonari, M. K.

Chimpanzee sample preparation: H.I, K. Nakabayashi.

Study conception: H. Kobayashi, M.C.L.

Funding: H. Kobayashi, K.K.

---

**Figure 1 - figure supplement 1.**
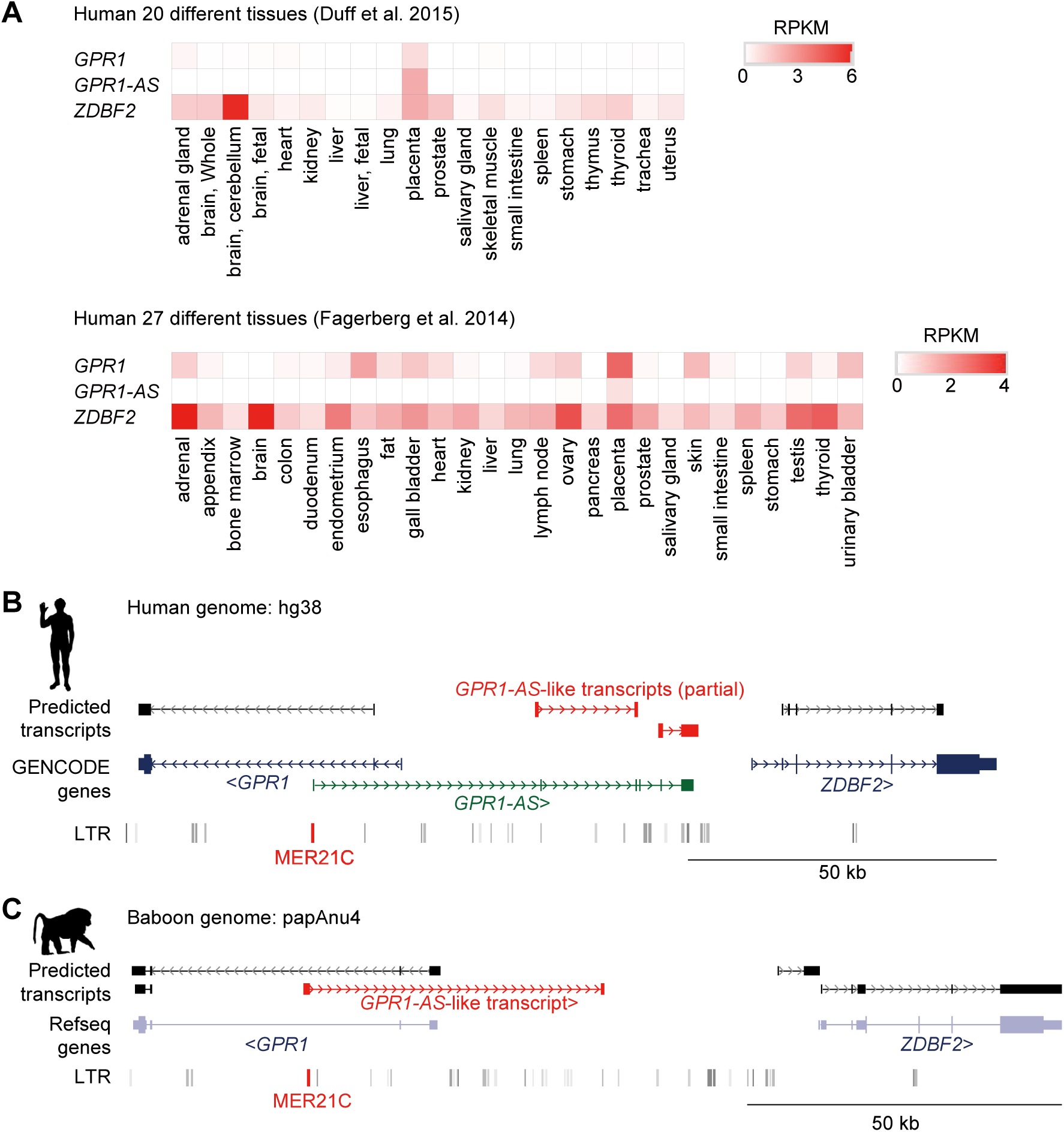
Identification of *GPR1-AS* orthologs using public and non-directional RNA-seq data. (A) Heat map showing the expression levels of *GPR1*, *GPR1-AS*, and *ZDBF2* in different human tissues, including the placenta. Genome browser screenshots of the GPR1- *ZDBF2* locus in humans (B) and baboons (C). Predicted transcripts were generated using public non-directional placental RNA-seq datasets (accession numbers: SRR1850957 for humans, GSM4696517 for baboons). Transcript/gene information and LTR retrotransposon positions are shown. *GPR1-AS*-like transcripts and MER21C retrotransposons are shown in red.

**Figure 2 – figure supplement 1.**
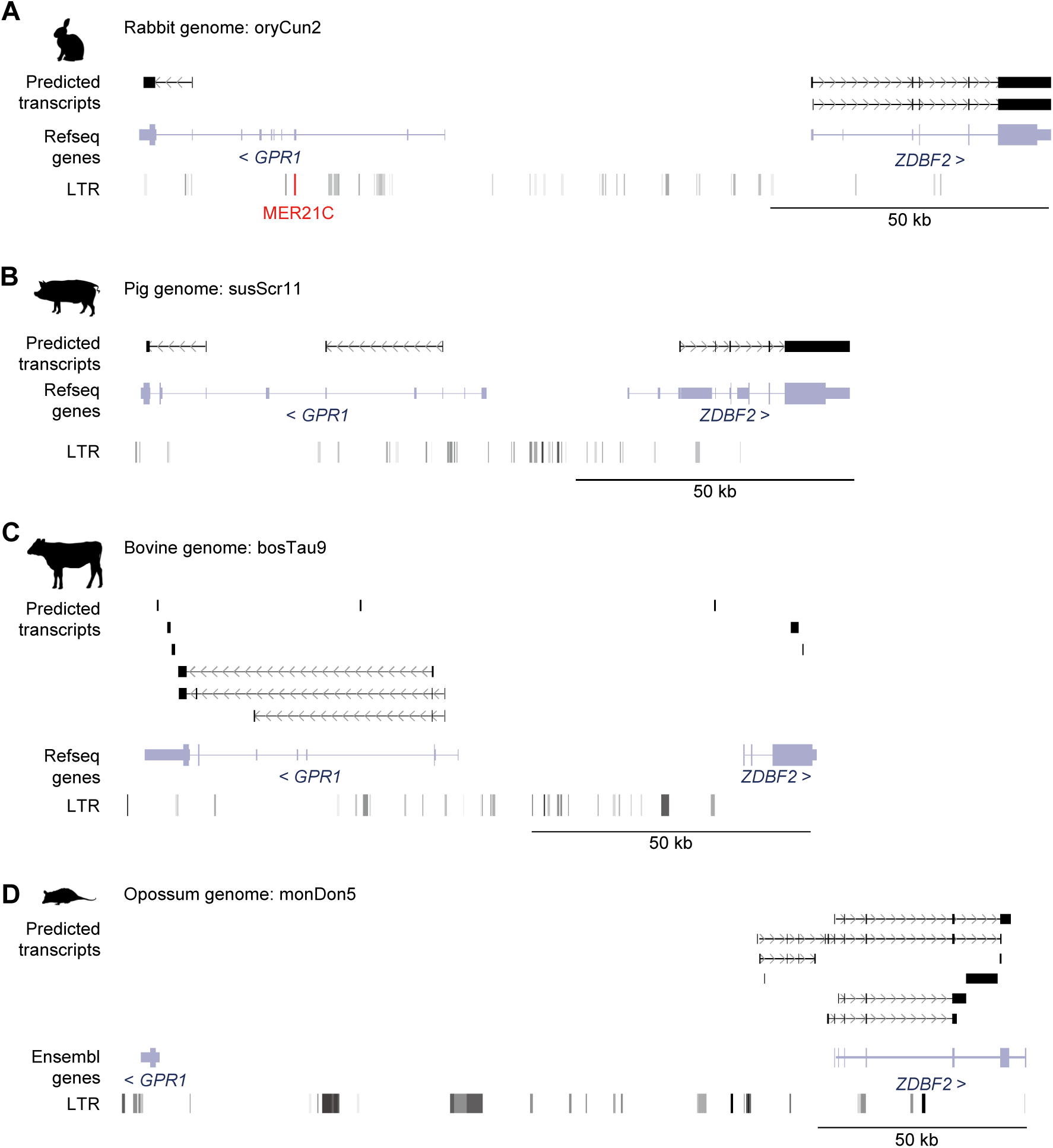
Search for *GPR1-AS* orthologs from embryonic transcriptomes. Predicted transcripts were generated using directional RNA-seq datasets of embryonic proper tissues from rabbit (A), pig (B), bovine (C), and opossum (D) embryos. Transcript/gene information and LTR retrotransposon positions are displayed and the annotated MER21C retrotransposon (only in rabbit) is highlighted in red.

**Figure 3 – figure supplement 1.**
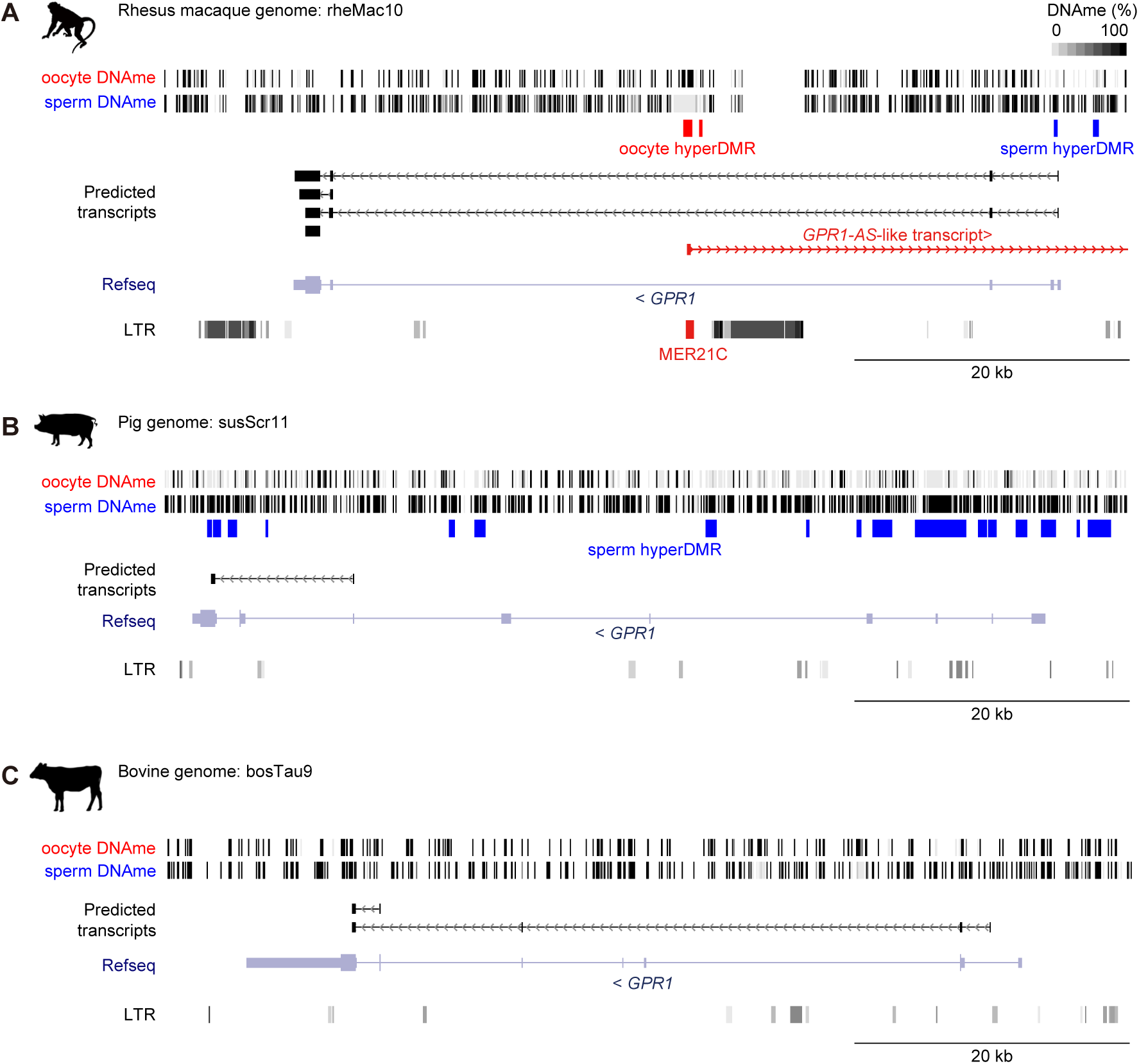
Search for germline DMRs from oocyte and sperm DNA methylomes. The DNA methylation (DNAme) levels of individual CpG sites in oocyte and sperm from rhesus macaque (**A**), pig (**B**), and bovine (**C**) whole genome bisulfite sequencing datasets are shown. Oocyte-methylated and sperm-methylated DMRs are highlighted in red and blue, respectively. Predicted transcripts from placental and extra-embryonic directional RNA-seq datasets (shown in Figure 1, **2**), genes annotated from RefSeq databases, and LTR positions from UCSC/RepeatMasker are included, with a MER21C retrotransposon overlapping rhesus macaque *GPR1-AS* highlighted in red.

**Figure 4 – figure supplement 1.**
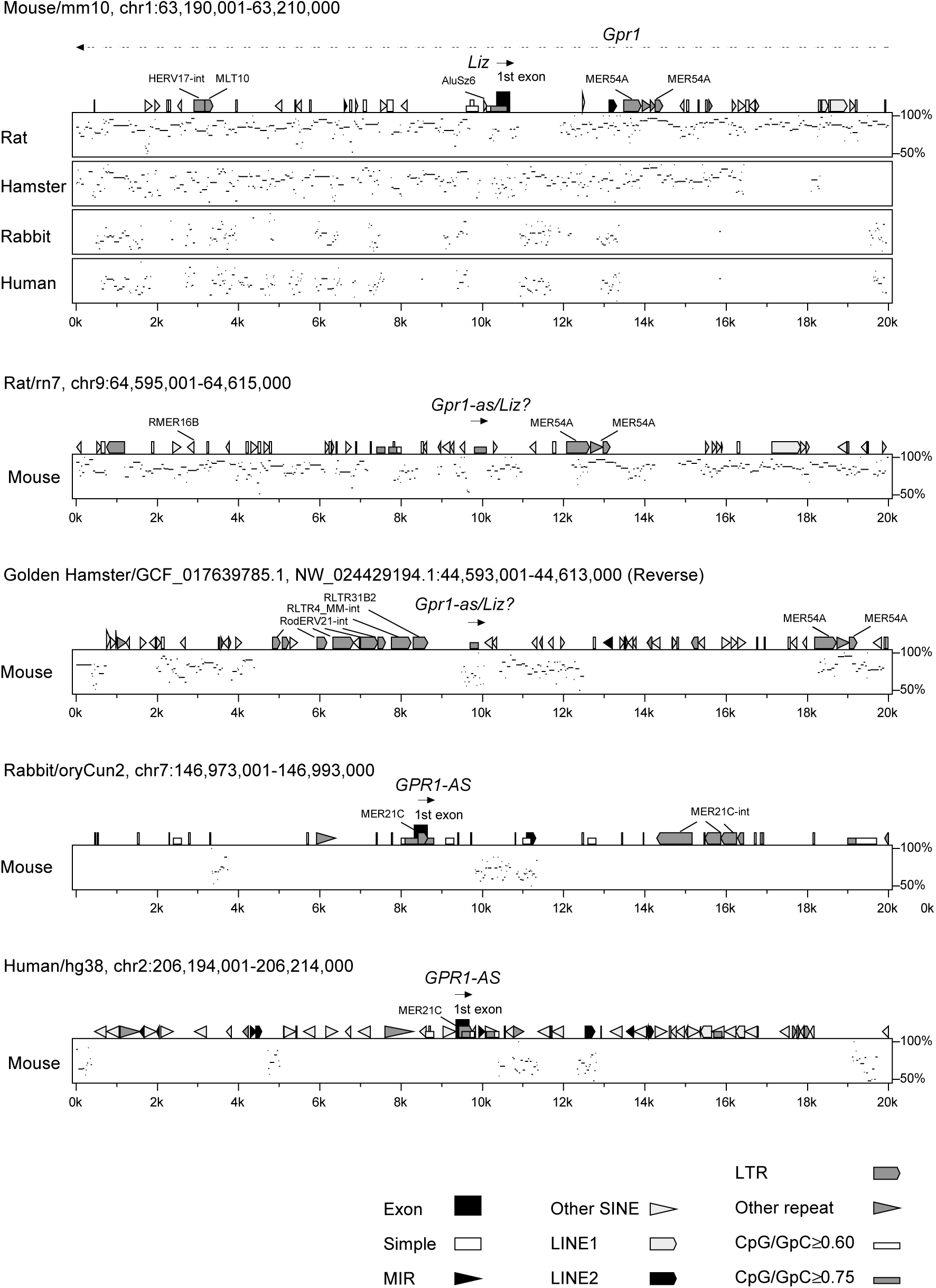
Reanalysis of repeat positions using RepeatMasker. Repetitive elements were re-identified in five mammalian species: mouse, rat, and hamster—where MER21C, which overlaps the first exon of human *GPR1-AS*, was not found in the homologous region—and rabbit and human, where it was detected. The Percent Identity Plot (PIP, showing a conservation scale between sequences from 50% to 100% on the y-axis) illustrates the order and alignment of the 20 kb region surrounding the *GPR1-AS* (*Liz*) transcription start site in each mammalian chromosome. Detected repeat elements are displayed above each plot. RepeatMasking was performed under less stringent settings, including switching search engines from RMblast to HMMER and adjusting speed/sensitivity settings from default to slow. Despite these adjustments, MER21C insertion was not detected in the three rodent species.

**Figure 4 – figure supplement 2.**
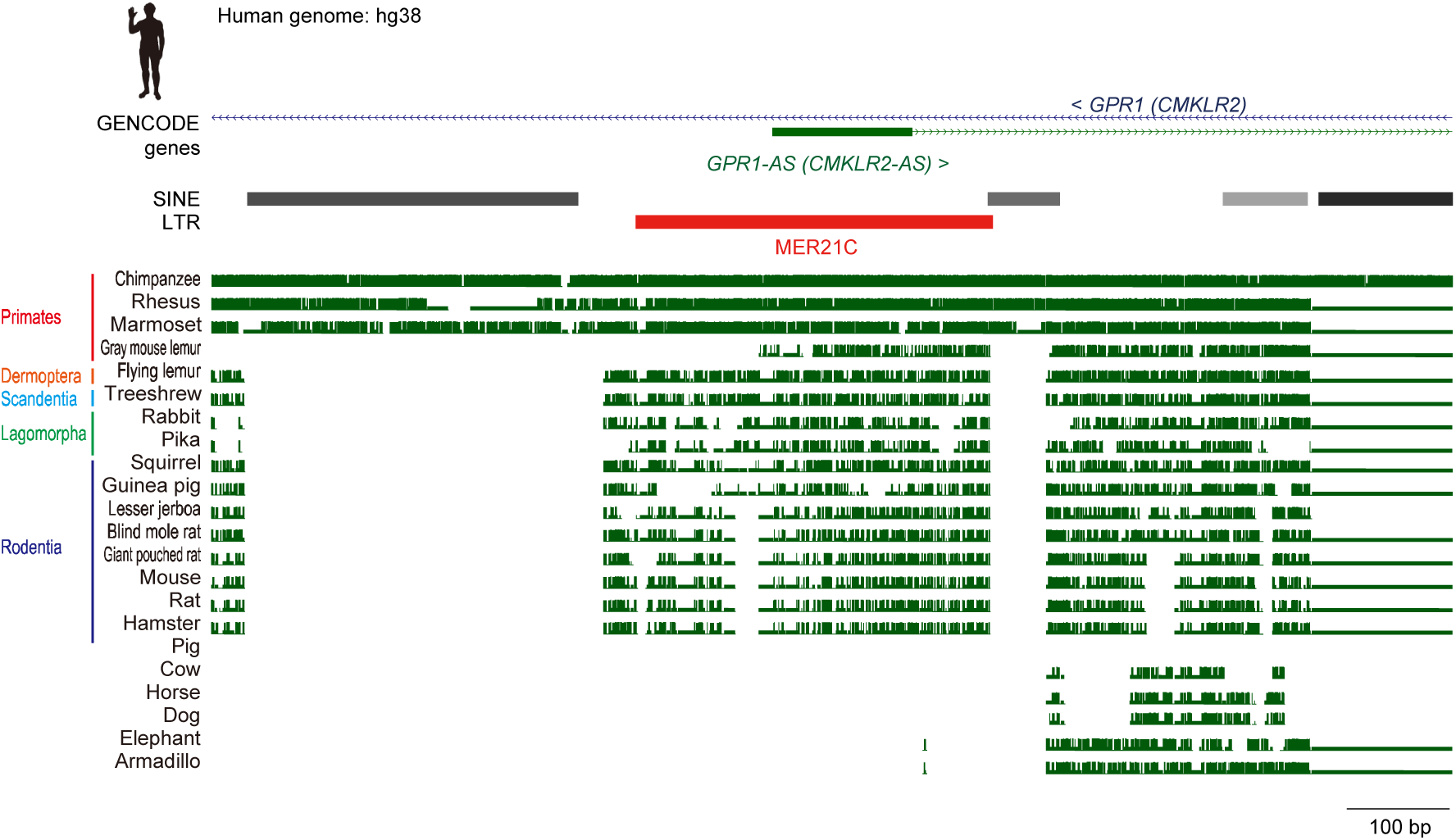
Multiple genome alignments at the first exon of *GPR1-AS* locus. Cactus generates reference-free, whole-genome multiple alignments (Armstrong et al. 2020). The Cactus track from UCSC Genome Browser displays multiple alignments across vertebrate species and evolutionary conservation metrics from the Zoonomia Project (Zoonomia 2020). Green square brackets indicate shorter alignments where DNA from one genomic context in the aligned species is nested within a larger alignment chain from a different genomic context. The alignment within these brackets may represent a short misalignment, a lineage-specific insertion of a retrotransposon in the human genome that aligns to a paralogous copy in another species. SINE and LTR retrotransposon positions from UCSC Genome Browser are also displayed.

**Figure 5 – figure supplement 1.**
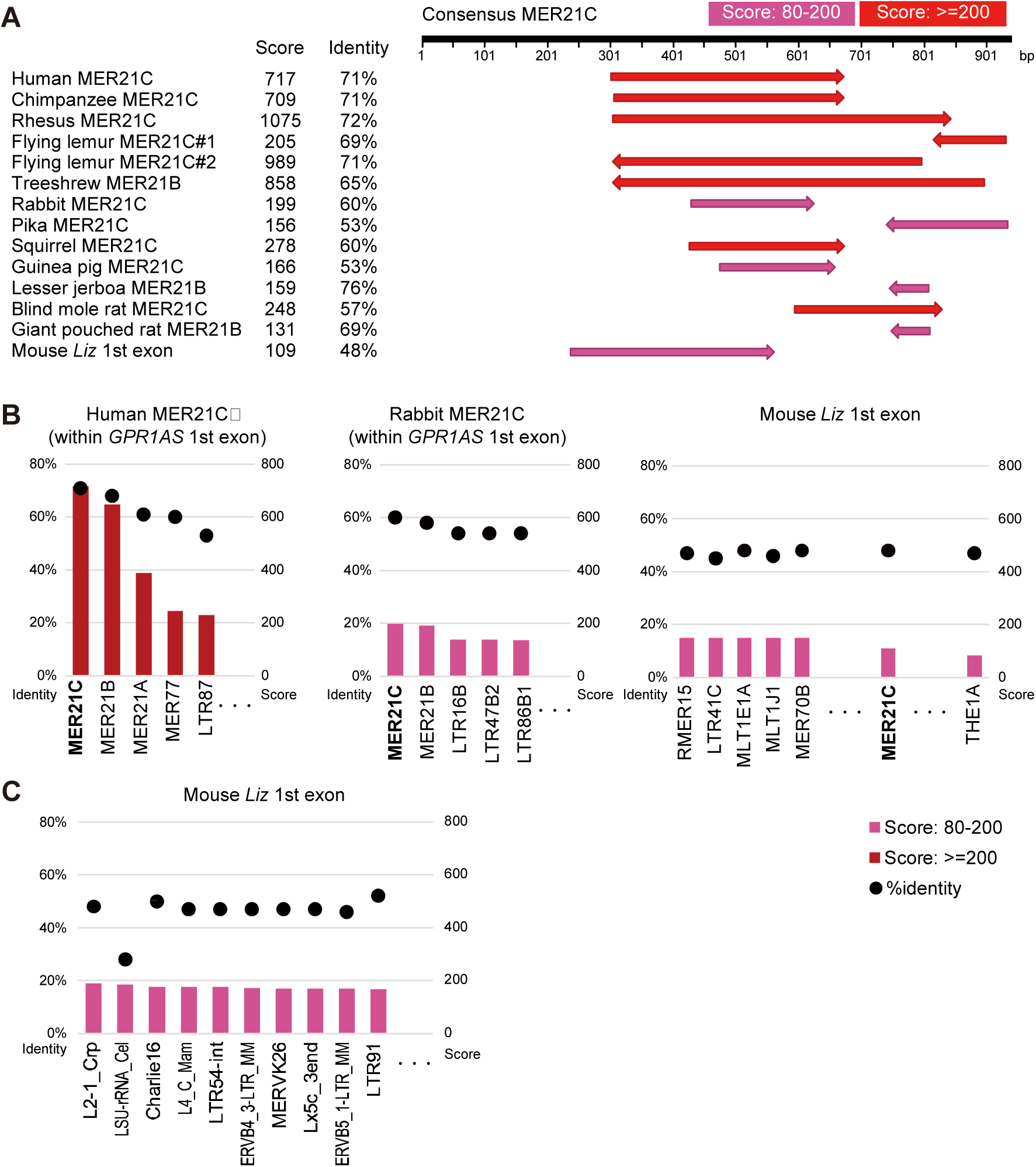
Pairwise alignment between consensus sequences of retrotransposons and *GPR1-AS*-exonic MER21 sequences. **(A)** MER21C (or MER21B) sequences located in the *GPR1* intron of eutherian genomes and the first exon of mouse *Liz* were compared with the consensus MER21C sequence. **(B)** Human and rabbit MER21C sequences overlapping the first exon of *GPR1-AS* and the first exon of mouse *Liz* were compared with the consensus sequences of ERV3/ERVL solo-LTRs present in human and mouse (n = 182). Each graph displays the identity percentages and alignment scores for the top five LTRs with the highest scores. In humans and rabbits, MER21C showed the highest identity with the exonic sequences. **(C)** The first exon of mouse *Liz* was compared with the consensus sequences of all retrotransposons present in mice (n = 1,361). The graph represents the top 10 retrotransposons with the highest scores. In mice, MER21C does not show sufficient sequence identity to the first exon of *Liz* to distinguish it from other retrotransposons. Pairwise alignment scores and percent identity values for each sequence pair were calculated using Genetyx software.

**Figure 5 – figure supplement 2.**
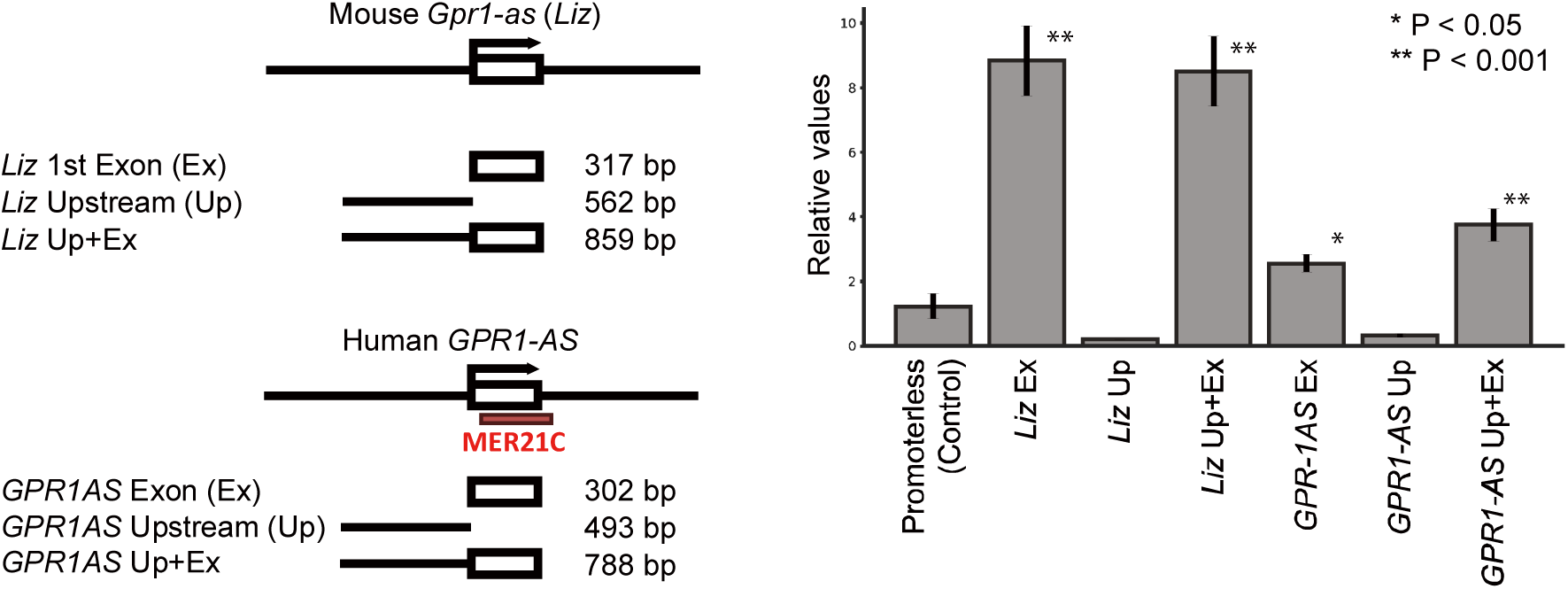
Promoter activities of first exons of mouse *Liz* and human *GPR1-AS*. (**A**) Constructs (inserted sequences) used for dual luciferase reporter assays in HEK293T cells. A promoter-less vector served as the negative control. (**B**) Results of dual luciferase reporter assays. Relative fold changes in Firefly luciferase activity (Firefly/Renilla) were normalized to the Firefly/Renilla ratio of the negative control. Error bars indicate mean ± s.e.m. Statistical significance was determined using unpaired t-tests: *P < 0.05, **P < 0.01. Data represent four biological replicates.

**Figure 5 – figure supplement 3.**
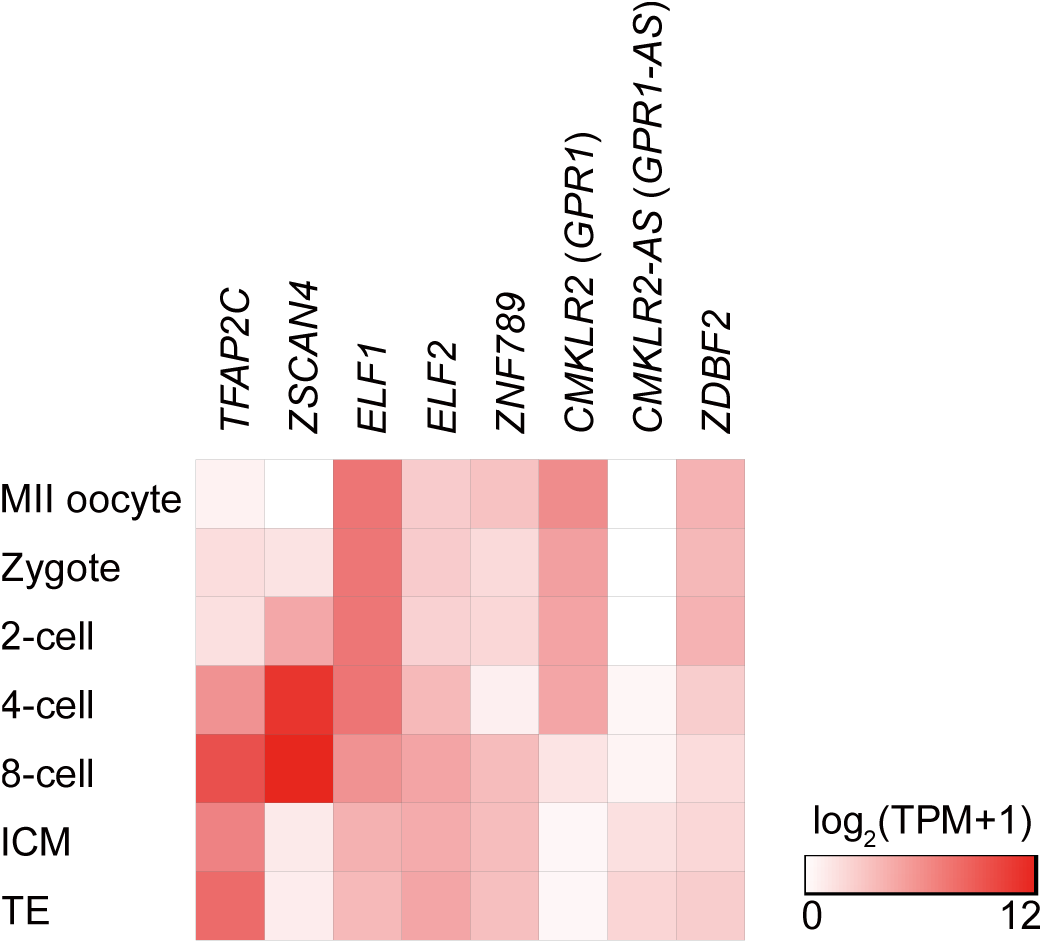
Expression patterns of transcription factors and imprinted genes during human preimplantation development. A heat map displaying the average expression of four transcription factors associated to human *GPR1-AS* or mouse *Liz* transcription, a primary KRAB-ZFP that binds to MER21C, and three imprinted genes surrounding the *ZDBF2* locus.

**Figure 6 – figure supplement 1.**
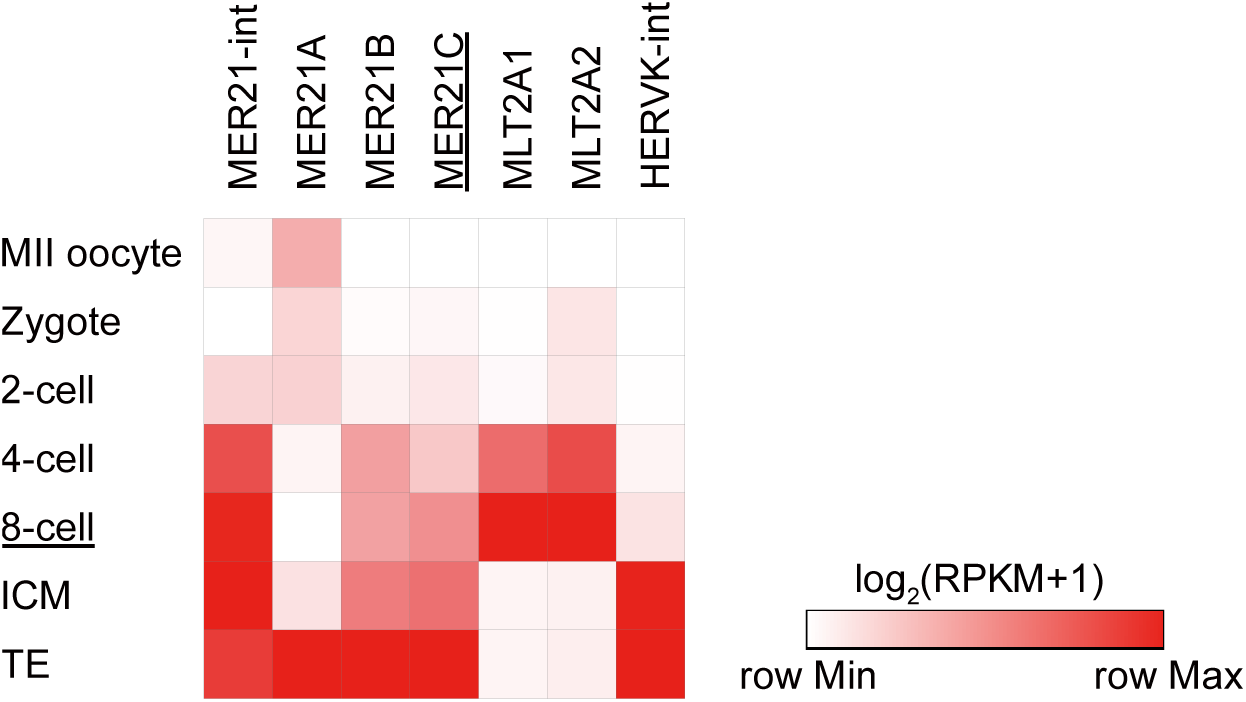
Human LTR reactivation during preimplantation development. Heat map displaying the average expression of select LTR retrotransposon families in human oocytes and early embryos. MLT2A1/MLT2A2 and HERVK are reactivated between the 4- to 8-cell stage and after the 8-cell stage, respectively (Grow et al. 2015; Hashimoto et al. 2021).

**Figure 7 – figure supplement 1.**
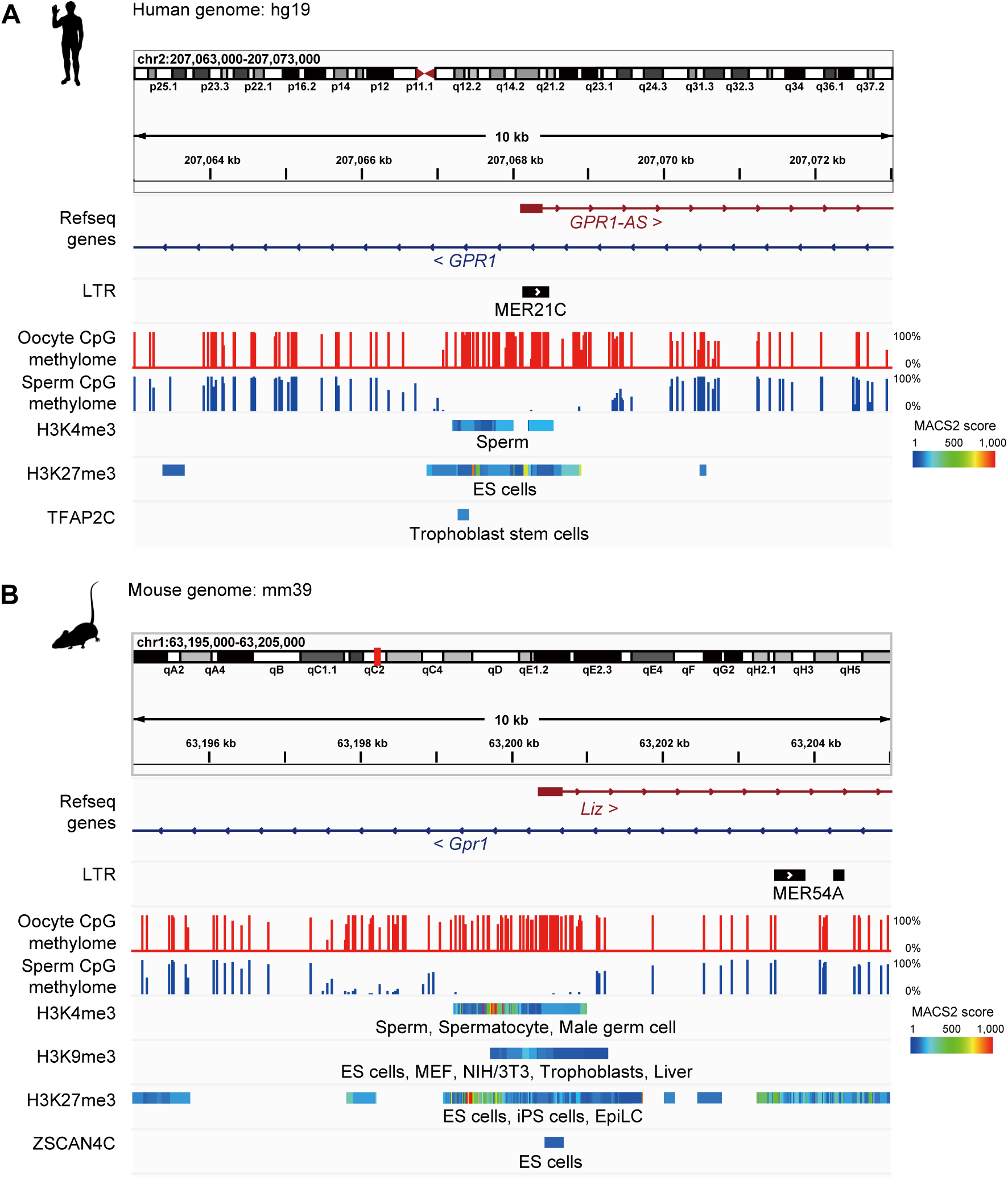
Interspecies epigenomic comparisons between human *GPR1-AS* and mouse *Liz*. IGV screenshots of the first exon of *GPR1-AS*/*Liz* in human (A) and mouse (B) showing DNA methylation, enrichment of post-translational histone modifications (H3K4me3, H3K9me3, and H3K27me3) and transcription factor binding sites (TFAP2C and ZSCAN4C) from ChIP-Atlas in various tissues. DNA methylomes from oocyte and sperm from mouse and human were published previously (Brind’Amour et al. 2018).

## REFERENCES

Akizawa H, Kobayashi K, Bai H, Takahashi M, Kagawa S, Nagatomo H, Kawahara M. 2018. Reciprocal regulation of TEAD4 and CCN2 for the trophectoderm development of the bovine blastocyst. Reproduction 155: 563–571.

Armstrong DL, McGowen MR, Weckle A, Pantham P, Caravas J, Agnew D, Benirschke K, Savage-Rumbaugh S, Nevo E, Kim CJ et al. 2017. The core transcriptome of mammalian placentas and the divergence of expression with placental shape. Placenta 57: 71–78.

Armstrong J, Hickey G, Diekhans M, Fiddes IT, Novak AM, Deran A, Fang Q, Xie D, Feng S, Stiller J et al. 2020. Progressive Cactus is a multiple-genome aligner for the thousand-genome era. Nature 587: 246–251.

Barau J, Teissandier A, Zamudio N, Roy S, Nalesso V, Herault Y, Guillou F, Bourc’his D. 2016. The DNA methyltransferase DNMT3C protects male germ cells from transposon activity. Science 354: 909–912.

Bogutz AB, Brind’Amour J, Kobayashi H, Jensen KN, Nakabayashi K, Imai H, Lorincz MC, Lefebvre L. 2019. Evolution of imprinting via lineage-specific insertion of retroviral promoters. Nat Commun 10: 5674.

Brind’Amour J, Kobayashi H, Richard Albert J, Shirane K, Sakashita A, Kamio A, Bogutz A, Koike T, Karimi MM, Lefebvre L et al. 2018. LTR retrotransposons transcribed in oocytes drive species-specific and heritable changes in DNA methylation. Nat Commun 9: 3331.

Chotalia M, Smallwood SA, Ruf N, Dawson C, Lucifero D, Frontera M, James K, Dean W, Kelsey G. 2009. Transcription is required for establishment of germline methylation marks at imprinted genes. Genes Dev 23: 105–117.

Duff MO, Olson S, Wei X, Garrett SC, Osman A, Bolisetty M, Plocik A, Celniker SE, Graveley BR. 2015. Genome-wide identification of zero nucleotide recursive splicing in Drosophila. Nature 521: 376–379.

Duffie R, Ajjan S, Greenberg MV, Zamudio N, Escamilla del Arenal M, Iranzo J, Okamoto I, Barbaux S, Fauque P, Bourc’his D. 2014. The Gpr1/Zdbf2 locus provides new paradigms for transient and dynamic genomic imprinting in mammals. Genes Dev **28**: 463-478.

Fagerberg L, Hallstrom BM, Oksvold P, Kampf C, Djureinovic D, Odeberg J, Habuka M, Tahmasebpoor S, Danielsson A, Edlund K et al. 2014. Analysis of the human tissue-specific expression by genome-wide integration of transcriptomics and antibody-based proteomics. Mol Cell Proteomics 13: 397–406.

Gao F, Niu Y, Sun YE, Lu H, Chen Y, Li S, Kang Y, Luo Y, Si C, Yu J et al. 2017. De novo DNA methylation during monkey pre-implantation embryogenesis. Cell Res 27: 526–539.

Glaser J, Iranzo J, Borensztein M, Marinucci M, Gualtieri A, Jouhanneau C, Teissandier A, Gaston-Massuet C, Bourc’his D. 2022. The imprinted Zdbf2 gene finely tunes control of feeding and growth in neonates. Elife 11.

Greenberg MV, Glaser J, Borsos M, Marjou FE, Walter M, Teissandier A, Bourc’his D. 2017. Transient transcription in the early embryo sets an epigenetic state that programs postnatal growth. Nat Genet 49: 110–118.

Grow EJ, Flynn RA, Chavez SL, Bayless NL, Wossidlo M, Wesche DJ, Martin L, Ware CB, Blish CA, Chang HY et al. 2015. Intrinsic retroviral reactivation in human preimplantation embryos and pluripotent cells. Nature 522: 221–225.

Hanna CW, Perez-Palacios R, Gahurova L, Schubert M, Krueger F, Biggins L, Andrews S, Colome-Tatche M, Bourc’his D, Dean W et al. 2019. Endogenous retroviral insertions drive non-canonical imprinting in extra-embryonic tissues. Genome Biol 20: 225.

Hashimoto K, Jouhilahti EM, Tohonen V, Carninci P, Kere J, Katayama S. 2021. Embryonic LTR retrotransposons supply promoter modules to somatic tissues. Genome Res 31: 1983–1993.

Hiura H, Sugawara A, Ogawa H, John RM, Miyauchi N, Miyanari Y, Horiike T, Li Y, Yaegashi N, Sasaki H et al. 2010. A tripartite paternally methylated region within the Gpr1-Zdbf2 imprinted domain on mouse chromosome 1 identified by meDIP- on-chip. Nucleic Acids Res 38: 4929–4945.

Huttley GA, Wakefield MJ, Easteal S. 2007. Rates of genome evolution and branching order from whole genome analysis. Mol Biol Evol 24: 1722–1730.

Imbeault M, Helleboid PY, Trono D. 2017. KRAB zinc-finger proteins contribute to the evolution of gene regulatory networks. Nature 543: 550–554.

Ivanova E, Canovas S, Garcia-Martinez S, Romar R, Lopes JS, Rizos D, Sanchez-Calabuig MJ, Krueger F, Andrews S, Perez-Sanz F et al. 2020. DNA methylation changes during preimplantation development reveal inter-species differences and reprogramming events at imprinted genes. Clin Epigenetics 12: 64.

Kai Y, Mei H, Kawano H, Nakajima N, Takai A, Kumon M, Inoue A, Yamashita N. 2022. Transcriptomic signatures in trophectoderm and inner cell mass of human blastocysts classified according to developmental potential, maternal age and morphology. PLoS One 17: e0278663.

Kaneko-Ishino T, Ishino F. 2012. The role of genes domesticated from LTR retrotransposons and retroviruses in mammals. Front Microbiol 3: 262.

Kaneko-Ishino T, Ishino F. 2022. The Evolutionary Advantage in Mammals of the Complementary Monoallelic Expression Mechanism of Genomic Imprinting and Its Emergence From a Defense Against the Insertion Into the Host Genome. Front Genet 13: 832983.

Karimi MM, Goyal P, Maksakova IA, Bilenky M, Leung D, Tang JX, Shinkai Y, Mager DL, Jones S, Hirst M et al. 2011. DNA methylation and SETDB1/H3K9me3 regulate predominantly distinct sets of genes, retroelements, and chimeric transcripts in mESCs. Cell Stem Cell 8: 676–687.

Kiyonari H, Kaneko M, Abe T, Shiraishi A, Yoshimi R, Inoue KI, Furuta Y. 2021. Targeted gene disruption in a marsupial, Monodelphis domestica, by CRISPR/Cas9 genome editing. Curr Biol **31**: 3956-3963 e3954.

Kobayashi H. 2021. Canonical and Non-canonical Genomic Imprinting in Rodents. Front Cell Dev Biol 9: 713878.

Kobayashi H, Sakurai T, Imai M, Takahashi N, Fukuda A, Yayoi O, Sato S, Nakabayashi K, Hata K, Sotomaru Y et al. 2012a. Contribution of intragenic DNA methylation in mouse gametic DNA methylomes to establish oocyte-specific heritable marks. PLoS Genet 8: e1002440.

Kobayashi H, Sakurai T, Sato S, Nakabayashi K, Hata K, Kono T. 2012b. Imprinted DNA methylation reprogramming during early mouse embryogenesis at the Gpr1- Zdbf2 locus is linked to long cis-intergenic transcription. FEBS Lett 586: 827–833.

Kobayashi H, Yamada K, Morita S, Hiura H, Fukuda A, Kagami M, Ogata T, Hata K, Sotomaru Y, Kono T. 2009. Identification of the mouse paternally expressed imprinted gene Zdbf2 on chromosome 1 and its imprinted human homolog ZDBF2 on chromosome 2. Genomics 93: 461–472.

Kobayashi H, Yanagisawa E, Sakashita A, Sugawara N, Kumakura S, Ogawa H, Akutsu H, Hata K, Nakabayashi K, Kono T. 2013. Epigenetic and transcriptional features of the novel human imprinted lncRNA GPR1AS suggest it is a functional ortholog to mouse Zdbf2linc. Epigenetics 8: 635–645.

Matsui T, Leung D, Miyashita H, Maksakova IA, Miyachi H, Kimura H, Tachibana M, Lorincz MC, Shinkai Y. 2010. Proviral silencing in embryonic stem cells requires the histone methyltransferase ESET. Nature 464: 927–931.

Matsushima W, Planet E, Trono D. 2024. Ancestral genome reconstruction enhances transposable element annotation by identifying degenerate integrants. Cell Genom 4: 100497.

Mei H, Kozuka C, Hayashi R, Kumon M, Koseki H, Inoue A. 2021. H2AK119ub1 guides maternal inheritance and zygotic deposition of H3K27me3 in mouse embryos. Nat Genet 53: 539–550.

Mika K, Whittington CM, McAllan BM, Lynch VJ. 2022. Gene expression phylogenies and ancestral transcriptome reconstruction resolves major transitions in the origins of pregnancy. Elife 11.

Moore T, Haig D. 1991. Genomic imprinting in mammalian development: a parental tug-of-war. Trends Genet 7: 45–49.

Morcos L, Ge B, Koka V, Lam KC, Pokholok DK, Gunderson KL, Montpetit A, Verlaan DJ, Pastinen T. 2011. Genome-wide assessment of imprinted expression in human cells. Genome Biol 12: R25.

Necsulea A, Soumillon M, Warnefors M, Liechti A, Daish T, Zeller U, Baker JC, Grutzner F, Kaessmann H. 2014. The evolution of lncRNA repertoires and expression patterns in tetrapods. Nature 505: 635–640.

Nicholls RD. 2000. The impact of genomic imprinting for neurobehavioral and developmental disorders. J Clin Invest 105: 413–418.

Papuchova H, Latos PA. 2022. Transcription factor networks in trophoblast development. Cell Mol Life Sci 79: 337.

Richard Albert J, Kobayashi T, Inoue A, Monteagudo-Sanchez A, Kumamoto S, Takashima T, Miura A, Oikawa M, Miura F, Takada S et al. 2023. Conservation and divergence of canonical and non-canonical imprinting in murids. Genome Biol 24: 48.

Rosenkrantz JL, Gaffney JE, Roberts VHJ, Carbone L, Chavez SL. 2021. Transcriptomic analysis of primate placentas and novel rhesus trophoblast cell lines informs investigations of human placentation. BMC Biol 19: 127.

Saito S, Akizawa H, Furukawa E, Yanagawa Y, Bai H, Takahashi M, Kawahara M. 2022. Generation of viable calves derived from developmentally mature blastocysts produced by on-gel culture. J Reprod Dev 68: 330–334.

Sakashita A, Kitano T, Ishizu H, Guo Y, Masuda H, Ariura M, Murano K, Siomi H. 2023. Transcription of MERVL retrotransposons is required for preimplantation embryo development. Nat Genet 55: 484–495.

Takahashi N, Coluccio A, Thorball CW, Planet E, Shi H, Offner S, Turelli P, Imbeault M, Ferguson-Smith AC, Trono D. 2019. ZNF445 is a primary regulator of genomic imprinting. Genes Dev 33: 49–54.

Tucci V, Isles AR, Kelsey G, Ferguson-Smith AC, Erice Imprinting G. 2019. Genomic Imprinting and Physiological Processes in Mammals. Cell 176: 952–965.

Wu CI, Li WH. 1985. Evidence for higher rates of nucleotide substitution in rodents than in man. Proc Natl Acad Sci U S A 82: 1741–1745.

Zhang W, Chen F, Chen R, Xie D, Yang J, Zhao X, Guo R, Zhang Y, Shen Y, Goke J et al. 2019. Zscan4c activates endogenous retrovirus MERVL and cleavage embryo genes. Nucleic Acids Res 47: 8485–8501.

Zoonomia C. 2020. A comparative genomics multitool for scientific discovery and conservation. Nature 587: 240–245.

Zou Z, Ohta T, Miura F, Oki S. 2022a. ChIP-Atlas 2021 update: a data-mining suite for exploring epigenomic landscapes by fully integrating ChIP-seq, ATAC-seq and Bisulfite-seq data. Nucleic Acids Res 50: W175–W182.

Zou Z, Zhang C, Wang Q, Hou Z, Xiong Z, Kong F, Wang Q, Song J, Liu B, Liu B et al. 2022b. Translatome and transcriptome co-profiling reveals a role of TPRXs in human zygotic genome activation. Science 378: abo7923.

